# Bowled over or over bowled? Age-related changes in the performance of bowlers in Test match cricket

**DOI:** 10.1101/2020.05.24.113191

**Authors:** Jack Thorley

## Abstract

Data from elite professional sports players provide a valuable source of information on human performance and ageing. Functional declines in performance have been investigated across a wide range of sporting disciplines that vary in their need for physical strength, endurance, cognitive ability and motor skills, but rarely have researchers considered other sources of heterogeneity that can exist among individuals. Using information on all male bowlers to have played Test match cricket since the early 1970s, I separated age-dependent variation in bowling performance at the population-level into within-individual and between-individual (cohort) changes. I found no evidence for senescence in bowling performance as measured via economy rate or wicket-taking ability, irrespective of the style of the bowler (fast or slow). Instead, analyses detected strong between-individual contributions to bowling performance as higher quality bowlers were able to compete at the elite level for longer, and were therefore over-represented in older age classes. Bowlers also experienced a deterioration in the last year of their Test careers. These results highlight that the very best Test match bowlers have been able to maintain and often improve their skill level well into their thirties, but how they accomplish this alongside the physical demands of Test cricket remains unresolved. Further multivariate models also identified a negative relationship among slow bowlers between their economy rate and their wicket-taking ability, suggesting that in general, the most economical slow bowlers in the modern era of Test match cricket have also taken wickets at the fastest rate. The same is not true for fast bowlers, which is perhaps partly because bowling at high speed compromises accuracy and thus increases scoring opportunities for batsman.

## INTRODUCTION

The vast majority of species experience declines in functional capacity in the later stages of their lifespan (Monaghan et al., 2008; Nussey et al., 2013; Jones et al., 2014; Shefferson et al., 2017). Contemporary human populations are no exception (Masoro and Austad, 2010), and the physiological deterioration that comes with old age (*senescence* or *ageing*) is perhaps better understood in humans than in any other species. Older humans have lower cognitive ability (Murman, 2015), respiratory capacity (Sharma and Goodwin, 2006), immune function (Pawelec, 2012), and muscle mass (Doherty, 2003) than younger humans, and together, such factors contribute to the increasing morbidity and mortality observed in the elder members of society worldwide (Uhlenberg, 2009; Vaupel, 2010). Yet despite often being associated with old age, declines in some physiological functions often begin much earlier in the life span than others (Walker and Herndon, 2010; see also Hayward et al., 2015 for example in a non-human population; Gaillard and Lemaître, 2017) and are not necessarily synonymous with debility. For example, some aspects of cognitive decline already begin in healthy, educated adults as early as their mid-to-late twenties (Salthouse, 2009), whereas the most pronounced declines in muscle wasting (sarcopenia) do not begin until individuals are over 50 years of age (Doherty, 2003). Understanding how and why the output of different physiological functions varies so markedly across human life span, and the consequences for health outcomes, remains a major challenge for modern gerontology (Christensen et al., 2009), but is complicated by the large variability introduced by differing lifestyles and the methodological difficulties of separating genetic, epigenetic, and environmental components of ageing (Christensen et al., 2006; Hjelmborg et al., 2006; Steves et al., 2012; Passarino et al., 2016).

Since the start of the 20^th^ century numerous studies have suggested that data from elite sportsmen and sportswomen can be particularly useful for investigating human ageing (Hill, 1925; Moore, 1975; Stones and Kozma, 1984; Careau and Wilson, 2017), as well as for testing predictions from evolutionary theory (Brooks et al., 2004; Pollet et al., 2013; Postma, 2014; Lailvaux, 2018). Using sporting data for these purposes bears the distinct advantage that elite sportsmen and women train rigorously throughout their sporting careers and show uniformly high motivation to succeed in their chosen discipline. As a result, analyses of age-related changes in performance in sportspeople reduce many of the biases surrounding differences in lifestyle and behaviour that are present among the general population. Furthermore, data collected in sporting competition is typically standardised; often provides large numbers of repeated records from individuals; and because additional extraneous factors are measured and recorded, they can be controlled for statistically.

The accumulated information from a large number of sports datasets converges upon the finding that peak physiological function in humans occurs on or around thirty years of age, and declines modestly thereafter, though the impact of age tends to depend upon the relative requirement for athleticism versus skill in the sport in question (Schulz and Curnow, 1988; Trappe, 2007; Bradbury, 2009; Baker and Tang, 2010). In general, sports involving greater speed and stamina, as exemplified by track and field events in athletics, favour younger individuals (Moore, 1975; Stones and Kozma, 1984; Young and Starkes, 2005), whereas those requiring more fine-control motor skills and less intense physical exertion tend to exhibit more delayed downturns (Baker et al., 2007; Schorer and Baker, 2009). This pattern is probably not surprising to any sports fan. The oldest ever male and female winners of 100m Olympic gold, Linford Christie and Fanny Blinkers-Koen, were 32 and 30 years of age when they won their medals in 1992 and 1948, respectively, and most winners have been considerably younger than this. If we contrast this to golf, a sport where success has historically been more dependent on motor skills than athletic ability, then we see that players have regularly competed on the world tour well into their fifties, and a number of male players have won the Masters beyond 40 years of age.

In the context of human performance, it is perhaps more illuminating to examine performance traits in sports demanding a more even contribution of skill and athleticism for success. Where this is the case, it is apparent that individuals can sometimes compensate for relative losses of function in physical attributes by increasing aspects of skill-based performance. Tennis and basketball provide two illustrative cases. Sutter et al. (2018) analysed first-serve performance in a large sample of modern professional tennis players and found that in both sexes, players experiencing more pronounced declines in average serve speed with increasing age displayed relative improvements in first serve accuracy. High average serve speed and high average accuracy were both associated with an increased probability of winning the match, but because both aspects of serving senesced, the compensatory increase in accuracy could only partly offset the overall performance declines in the later stages of player’s careers. In a separate study, Lailvaux et al. (2014) examined points-scoring ability in professional basketball players playing in the National Basketball Association (NBA). In male basketball players, but not females, the decline in points arising from ‘close-range’ two-point field throws began three to four years before that detected for ‘longer-range’ three-point field throws, which the authors argued is indicative of compensation: as declines in speed and power reduce the ability to score two-point field goals closer to the basket, players having the accuracy to continue scoring more distant three-point field goals can maintain points-scoring ability. Although further examples of such dynamic compensation in sports players are relatively few and far between (Schorer and Baker, 2009), we should not necessarily be surprised by their presence in humans as trait compensation is widespread across the animal kingdom (Husak et al., 2011; Cameron et al., 2013; Dennenmoser and Christy, 2013) and is underpinned by evolutionary theory grounded in trade-offs (Roff, 1992; Stearns, 1992).

Though sports datasets are often longitudinal in design, containing large numbers of repeated records from the same individuals, very few studies of sporting performance have explicitly sought to separate within-individual changes in performance from compositional differences between different age classes, or what are sometimes referred to as cohort effects. It is not that researchers in sports performance have failed to recognise cohort effects, for they have been discussed and applied in several places (Young and Starkes, 2005; Stones, 2019), but rather that they are rarely incorporated into analytical frameworks explicitly. There are two principal reasons why the inclusion of between-individual ‘cohort’ effects is warranted. Firstly, by failing to consider cohort effects, one risks under or overestimating age-related changes occurring within individuals and therefore fail to correctly quantify the ageing process. Secondly, by incorporating cohort effects one can gain important additional information about changes in human performance that would otherwise go undetected. For instance, if only the highest quality sports players enjoy long careers and compete professionally into the older age classes-because less competent performers have already stopped or are no longer up to the required standard-then this raises questions about how they are able to do so. Is their superiority gained earlier in their career and maintained? What aspects of their performance allow them to continue competing later than others? Do these long-lived professionals follow a different training regime? Equally, one might wonder whether professional sports players that start competing at the highest level at a very young age are relatively better or worse than individuals who started doing so later in life.

In this study, I examine age-related variation in the performance of bowlers in Test match cricket and dissect changes occurring within individuals from changes occurring between individuals. In so doing, I borrow methods from animal ecology where researchers routinely analyse long-term data taken from animals tracked across their life span to understand how individual variation in performance changes with age (van de Pol and Verhulst, 2006; Rebke et al., 2010; Nussey et al., 2011). Like animals in natural populations, cricketers at the elite international level are selected to perform to the best of their ability and their performance is measured relative to other individuals seeking to do so the same. They ‘appear’ in the population when they are deemed good enough to compete, have some scope to improve with experience, but should they fail to compete effectively, then they will be replaced and ‘disappear’ from the population. I focus on Test cricket as this provides the longest-running and most standardised form of the sport, often being considered the game’s highest standard. An abridged outline of cricket is provided in the methods, but in general, batsmen are specialised in the accruing of runs, while bowlers are concerned with dismissing (“getting out”) batters and reducing the rate at which they score runs (points). Bowlers can be broadly separated into fast bowlers, who bowl the ball quickly (110-150kph), and slow bowlers (70-110kph), who more often rely on guile and subtle variations in spin and trajectory to succeed. Both types of bowling require considerable motor skill and stamina (Noakes and Durandt, 2000; Petersen et al., 2010), though fast bowlers also tend to require a greater degree of athleticism in order to deliver the ball at speed from one end of the wicket to the other.

By analysing the complete dataset of modern era (here defined as post-1973) Test match bowlers, I find that two measures of bowling performance show no sign of senescent decline. I identify other important within-individual and among-individual contributions to bowling performance, and highlight a key difference in the relationship between wicket-taking ability and economy rate in fast and slow bowlers.

## MATERIALS AND METHODS

### a) “Test Cricket”

It would take too much space to detail all the nuances of cricket. For this, I recommend Woolmer’s *Art and Science of Cricket* (Woolmer, 2008). Instead, I provide enough information for readers to understand the basic principles by which each match proceeds, and the contributions that individuals make to team performance.

Test cricket is the longest format of cricket and is often considered the game’s highest standard. Since the first official ‘Test match’ was contested between England and Australia in 1877, 12 international teams have been granted Test status by the International Cricket Council and regularly compete against one another. As in all forms of cricket, either side fields eleven players composed of batters and bowlers.

Cricket matches are played in innings. In a single team innings, one team bats while the other fields. Test matches ordinarily comprise four innings with each team batting and bowling twice in alternation. The most basic aspect of play in a game of cricket is a “ball”, where a bowler, from the fielding team, propels a hard leather ball towards the batter attempting to get the batter “out”, while the batter attempts to hit the ball to score “runs” (points). A bowler can dismiss a batter by various means and thereby take their wicket at which point the batter leaves the field of play and is replaced by another member of the batting team. Batters may also be dismissed by the actions of other members of the fielding team independently of the bowler’s actions. Bowlers bowl to batsmen in six-ball periods known as an “over”. While a single bowler cannot bowl successive overs, there is no limit in Test cricket on the total number of overs a bowler can bowl within an innings. i.e. a bowler could bowl 25 overs in an innings should their captain wish them to do so, which would equate to 150 balls. Once the batting team loses 10 wickets (their innings is completed) the combined scores of all the batters generates a team total that the opposition team then aims to meet or surpass. Note that an individual batting performance is also called an innings.

Over a completed match, a team wins when they score more runs across their two innings than the opposition while also taking all of the opposition’s wickets. A draw occurs when one team scores more runs than the opposition without taking all of their wickets within the designated playing time. To try and force a win, the team that is batting may therefore decide to forfeit their remaining wickets (“declare”) in order to provide enough time for them to bowl out the opposition.

Because Test matches historically varied in length from three days to more than a week, the data was standardised by focussing only on cricketers who debuted after 14^th^ February 1973, from which point Test matches were uniformly restricted to a maximum of five days in length with a minimum of 6 hours of play a day, weather permitting. In so doing, the analyses were also restricted to modern cricketers who can be considered as elite sportsmen playing in an era when the sport was largely professional.

### b) The dataset

The bowling data of all male Test cricketers who debuted after 14^th^ February 1973 was extracted from *ESPN Cricinfo* (https://www.espncricinfo.com/). Specifically, the *Octoparse* software (https://www.octoparse.com/) was used to acquire the unique ESPN profile identifier of all individuals, as well as their date of birth and their bowling style. With the profile identifiers the complete bowling record of every individual could then be accessed from *Cricinfo* using the *getPlayerData()* function in the *cricketr* package (Ganesh, 2019) in R v 3.6.1 (Team, 2019). All data extraction was conducted on the 21^st^ June 2019. Bowlers were defined as cricketers who fielded at least 10 times in their Test career and bowled in at least 70 percent of all innings they fielded in. In order to correctly partition within- and between-individual effects, only individuals whose Test career was thought to have finished at the point of data extraction were considered, producing a dataset of 354 individuals (n = 16,279 per-individual innings bowled in). Bowlers were described as either fast bowlers (n = 227; including fast, fast-medium, and medium-fast bowlers) or slow bowlers (n = 128; medium-slow, and spinners) according to information on *Cricinfo* and *Wikipedia* so that any possible influence of bowling speed/style on age-related changes in performance could be separated. Spin bowlers represented 95.3% of the individuals in the dataset of slow bowlers.

### c) Univariate modelling of performance

To examine age-related variation in the performance of Test match bowlers, I fitted a series of univariate mixed-effects models using the within-group centering approach of Van de Pol and Verhulst (2006). This method separates age-related changes that take place within individuals (due to improvement or senescence) from changes that place between individuals at the population-level, as arise when individuals of differing quality appear or disappear from the population sample more or less often at different ages. Fast bowlers and slow bowlers were modelled separately, and in each case three performance attributes in each innings bowled were assessed, i-economy rate: the number of runs a bowler concedes per over, ii-wicket taking ability: the number of wickets taken by a bowler in an innings, and iii-proportion of the total overs of an team’s innings bowled by an individual. Economy rate was fitted to a Gaussian error distribution, wicket-taking ability to a negative binomial distribution with a zero-inflation parameter applied across all observations (i.e. zi ~ 1), and proportional overs bowled to a binomial distribution (logit link) with the number of overs bowled by an individual set as the numerator, and the total overs bowled in the innings set as the denominator. For all three attributes I first fitted a baseline model that included fixed effects of match innings number (four-level factor: 1,2,3,4), and whether the match was played at home or away (two-level factor). For the wicket-taking models I also included a term for the number of overs bowled by an individual. In controlling for the number of overs bowled, wicket-taking ability is thus largely synonymous with ‘strike rate’.

The baseline model was then compared to additional models incorporating different combinations of age-dependent or age-independent variables (Table 1). In these models, age was fitted as either a linear or quadratic function, a longevity term specifying the age of last Test match accounted for possible selective disappearance of poorer quality bowlers, and a factor denoting if it was the last year of a bowler’s career controlled for possible terminal declines in performance. Lastly, I specified further models that interacted the age terms with the innings number and the age of last appearance. As random effects I included random intercepts for the individual bowler, for the interaction between the country a bowler played for and the decade of the match, and for the interaction between the opposition and the decade of the match. The latter two effects controlled for differences in the quality of the Test-playing nations across time. In the economy rate models and the proportion of overs bowled models a random intercept of match was also included. Likelihood ratio testing found that the exclusion of the match effect in the wicket-taking models was preferable to a model in which the effect was present, which is probably because the number of wickets in a match (maximum 20 per team) is constrained in such a way that individuals that take a lot of wickets prevent others from doing so. Models were ranked on their Akaike Information Criterion (AIC) under maximum likelihood settings, and with ΔAIC < 2 were considered to show competing support (Burnham and Anderson, 2002). Where two models fell within 2 AIC units, the results of the simpler model are presented. All continuous variables were standardised to the mean and unit variance prior model fitting. Univariate models were fitted in the *lme4* and the *glmmTMB* R packages (Bates et al., 2015; Brooks et al., 2017).

**Table 1:** A comparison of models examining age-dependent and age-independent variation in the bowling performance of Test match cricketers across their careers. Separate models were fitted to fast bowlers and slow bowlers, for three performance metrics, economy rate (number of runs conceded per over per innings), wicket-taking ability (the number of wickets taken per innings), and the proportion of the overs in the innings bowled. Baseline refers to the baseline model as outlined in the main text. ALA-Age at last appearance; LY-Last year of career. The most parsimonious model in each case is highlighted in bold.

| Fast bowling |  | Economy Rate |  | Wicket-taking ability |  | Prop. innings overs bowled |
| --- | --- | --- | --- | --- | --- | --- |
| Model | k | ΔAIC | k | ΔAIC | k | ΔAIC |
| Baseline | 10 | 73.93 | 11 | 34.99 | 9 | 106.56 |
| Baseline + Age | 11 | 29.34 | 12 | 36.97 | 10 | 108.56 |
| Baseline + Age + Age<sup>2</sup> | 12 | 31.18 | 13 | 37.95 | 11 | 73.83 |
| Baseline + ALA + LY | 12 | 44.69 | 13 | 4.48 | 11 | 41.71 |
| Baseline + Age + ALA + LY | 13 | 6.46 | 14 | 5.83 | 12 | 39.69 |
| Baseline + Age + Age<sup>2</sup> + ALA + LY | 14 | 8.41 | 15 | 7.67 | 13 | 17.47 |
| Baseline + Age*Inns + ALA + LY | 16 | 0.00 | 17 | 0.00 | 15 | 37.85 |
| Baseline + Age*ALA + ALA + LY | 14 | 7.93 | 15 | 7.66 | 13 | 41.19 |
| Baseline + Age*Inns + Age*ALA + ALA + LY | 17 | 1.34 | 18 | 1.90 | 16 | 38.24 |
| Baseline + Age*Inns + Age<sup>2</sup>*Inns + ALA + LY | 20 | 6.67 | 21 | 1.99 | 19 | 19.52 |
| Baseline + Age*ALA + Age<sup>2</sup>*ALA + ALA + LY | 16 | 6.78 | 17 | 35.57 | 15 | 0.00 |
| Baseline + Age*Inns + Age<sup>2</sup>*Inns + Age*ALA + Age<sup>2</sup>*ALA + ALA + LY | 22 | 4.68 | 23 | 4.72 | 21 | 2.17 |
| Slow bowling |  | Economy Rate |  | Wicket-taking ability |  | Prop. innings overs bowled |
| Model | k | ΔAIC | k | ΔAIC | k | ΔAIC |
| Baseline | 10 | 18.22 | 11 | 20.70 | 9 | 57.14 |
| Baseline + Age | 11 | 17.51 | 12 | 22.66 | 10 | 58.26 |
| Baseline + Age + Age<sup>2</sup> | 12 | 16.98 | 13 | 24.57 | 11 | 54.52 |
| Baseline + ALA + LY | 12 | 0.00 | 13 | 1.15 | 11 | 36.74 |
| Baseline + Age + ALA + LY | 13 | 1.57 | 14 | 0.74 | 12 | 33.72 |
| Baseline + Age + Age<sup>2</sup> + ALA + LY | 14 | 1.46 | 15 | 14.10 | 13 | 33.98 |
| Baseline + Age*Inns + ALA + LY | 16 | 5.67 | 17 | 3.09 | 15 | 21.96 |
| Baseline + Age*ALA + ALA + LY | 14 | 1.76 | 15 | 0.00 | 13 | 28.74 |
| Baseline + Age*Inns + Age*ALA + ALA + LY | 17 | 5.90 | 18 | 2.33 | 16 | 17.02 |
| Baseline + Age*Inns + Age<sup>2</sup>*Inns + ALA + LY | 20 | 8.66 | 21 | 8.46 | 19 | 25.56 |
| Baseline + Age*ALA + Age<sup>2</sup>*ALA + ALA + LY | 16 | 4.26 | 17 | 2.62 | 15 | 8.48 |
| Baseline + Age*Inns + Age<sup>2</sup>*Inns + Age*ALA + Age<sup>2</sup>*ALA + ALA + LY | 22 | 11.86 | 23 | 8.65 | 21 | 0.00 |

### d) Multivariate bowlers

To investigate how wicket taking ability and economy rate covaried among individuals across their career, I fitted bivariate mixed models that included the performance metrics as a negative-binomial and Gaussian distributed response, respectively. The advantage of the multivariate approach is that it allows for the estimation of the posterior correlation between traits of interest after controlling for the confounding effects of other covariates (Houslay et al., 2018). Here, I was interested in the posterior among-individual correlation between the two performance traits. Fast and slow bowlers were again modelled separately, first with a model including only group-level (‘random’) effects, before a second model then included appropriate population-level ‘fixed’ effects, as informed by the best-fitting univariate models. The only exception to this was the exclusion of the age of last appearance term so as not to devalue individuals who played for a long time; so that for estimate of the posterior correlation any variance previously attributed to the longevity term was instead credited to the individual bowler and reflected in his individual intercept (see Tables S4 and S5 for full model table). Models were fitted in the *brms* package (Bürkner, 2018), with three chains of 3,000 iterations, of which 1,000 were dedicated to the warm-up. Posterior predictive checks highlighted adequate mixing of chains and appropriate use of default priors.

Lastly, the latter two multivariate models were refitted to two data sets that included cricketers that were still playing at the time of data gathering, or who only stopped playing Test cricket very recently. The updated fast bowler data set contained 274 individuals (n = 14033 per-individual innings), the slow bowler data set 160 individuals (n = 6413 total per-individual innings). This analysis was partly to estimate the posterior correlations with more data, but it also enabled the identification and ranking of players still playing Test cricket against the very best bowlers of the modern-era. For these models I also removed the population-level terminal effect as at the time of data extraction and analysis it was not possible to know whether current players were in the last year of their Test match career.

## RESULTS

### a) Age-related changes in cricketing performance

Patterns of age-related change in the performance of Test match bowlers varied according to bowling style, the innings of the match, and the performance metric that was considered (Figure 1, Figure 2, Table 1). In no case did models suggest that individuals were senescing in their economy rate, or in their wicket-taking ability. For fast bowlers, the best fitting model indicated that the economy rate of individuals improved (declined) with age (Figure 1a, Table S1). This decline in economy rate was evident in all innings (Table S1), but the slope tended towards being more significantly negative in the fourth innings of matches compared to the first three innings (p value for pairwise contrast between first, second, and thirds innings versus the fourth innings: 0.021, 0.011, and 0.068 respectively, Figure 2a).

**Figure 1.**
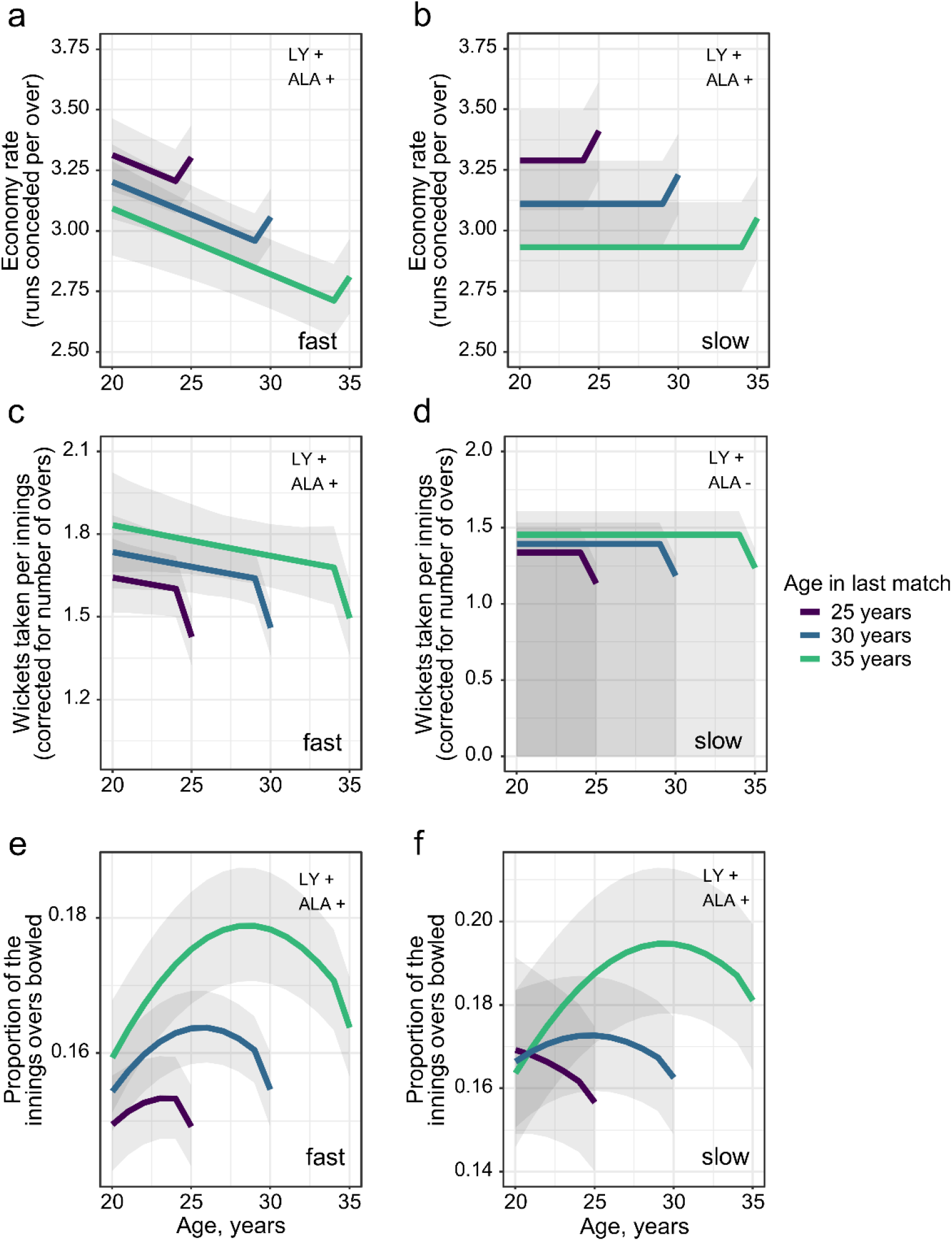
Age-related changes in the performance of Test match bowlers, separated by bowling style. Solid lines display the predicted performance from the best supported mixed effects models in each case for fast and slow bowlers; for economy rate (a, b), wicket-taking ability (c, d), and the proportion of the total overs in an innings bowled (e, f). Lines are coloured according to the age at which individuals stop playing Test match cricket, here estimated at the ages of 25, 30 and 35 years. Shading displays the 95% confidence intervals conditional on the fixed effects. For the fast bowlers, predictions were made for innings 1 at home, while for the slow bowlers, predictions were made for innings 4 at home. Note that a lower economy rate indicates higher performance. The significance (α = 0.05) of the presented terminal decline (LY-last year of career) or longevity effect (ALA-age at last appearance) for each model are noted with a +/− sign within the respective panels.

**Figure 2.**
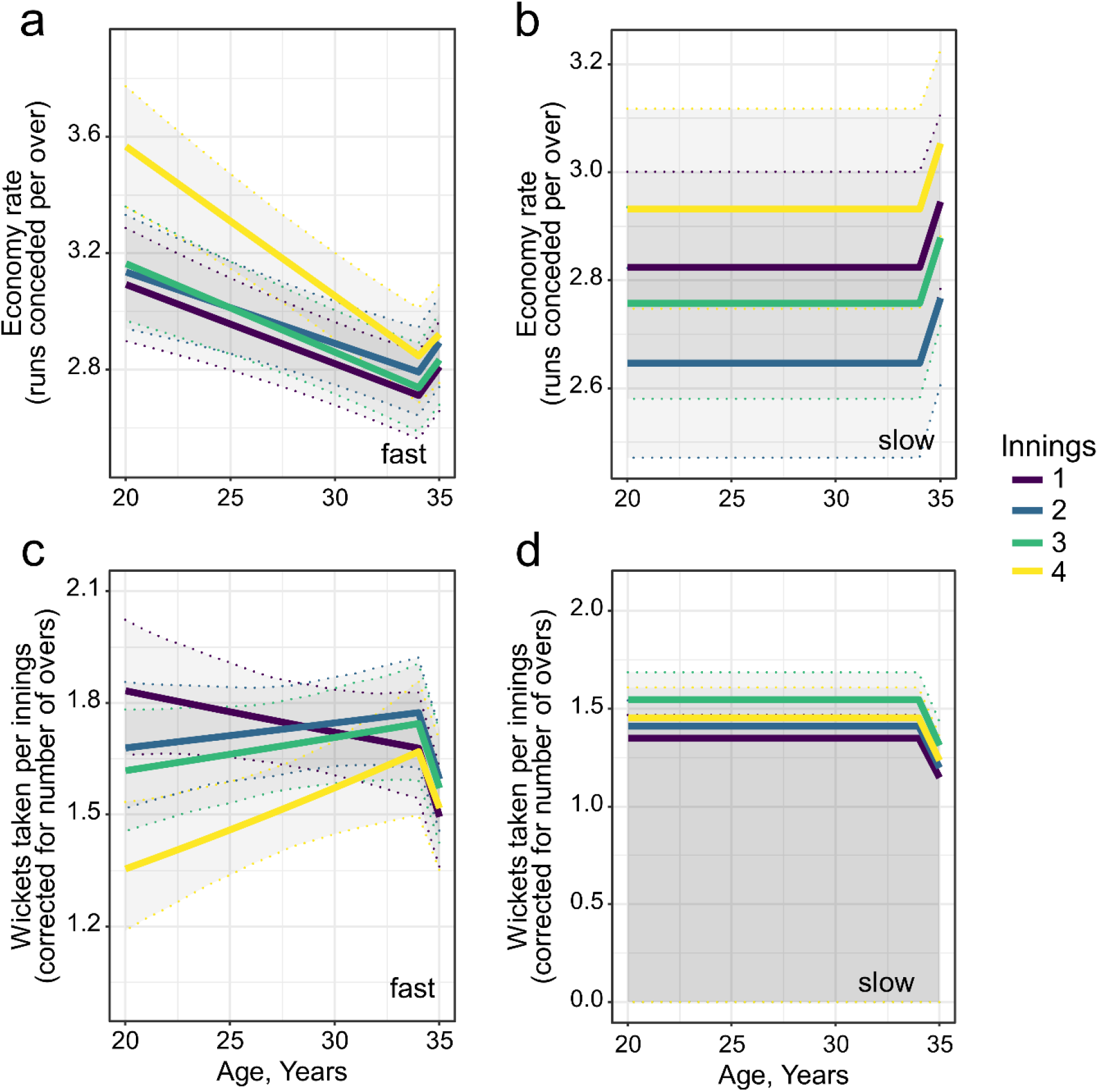
Age-related changes in the performance of Test match bowlers according to the innings of the match. Solid lines display the predicted performance from the best supported mixed effects models in each case for fast and slow bowlers, coloured according to the innings of the match; for economy rate (a, b), and wicket-taking ability (c, d). Shading displays the 95% confidence intervals conditional on the fixed effects. Predictions were made for individuals who played their last match when they were 35 years old, with a terminal decline in the last year of life included. Note that a lower economy rate indicates higher performance.

In general, the wicket-taking ability of fast bowlers also improved with age, though the magnitude of the change was dependent upon the innings in question (Table 1, Figure 2, Table S2). The overall wicket-taking ability of fast bowlers as indicated by the intercept term was graded across innings such that their success decreased from innings one to innings four (Figure 2). However, bowlers became more efficient at taking wickets in innings two, three, and four as they got older (Figure 2c). In contrast, the age effect in the first innings displayed a negative slope, though this effect was not significant (GLMM, fast bowlers estimate on log-link scale = − 0.024 ± 0.017, z-value = −1.46, p = 0.145). Taken together, these results imply that both the rate at which fast bowlers take wickets (the number of balls bowled per wicket or ‘strike rate’) and the number of they runs concede while taking those wickets improves across their career.

The economy rate and wicket-taking ability of slow bowlers followed a different trajectory to that of fast bowlers. After controlling for the strong effects of the innings number (LMM, χ^2^_3_ = 31.26, p < 0.001) and the location of the match (LMM, estimate away versus home = −0.169 ± 0.041, χ^2^_1_ = 16.86, p < 0.001) the economy rate of slow bowlers did not change with age (Figure 1, Figure 2, Table 1). The most parsimonious model for wicket-taking ability also suggested that wicket-taking ability was independent of age once the total number of overs bowled was controlled for (GLMM, estimate on log-link scale = 0.414 ± 0.014, z-value = 30.31, p < 0.001).

Both fast bowlers and slow bowlers bowled proportionally less of the total overs in an innings between 25 and 30 years of age (Table 1, Table S3). While this is indicative of reduced workload and could be related to physiological senescence, the size of the effect is small and equates to only minor reductions in the total workload of bowlers, at most 1-2%. At the mean innings duration in our dataset of 94.4 overs (1SD = 40.33), a 1% reduction in the number of overs bowled by a fast bowler represents roughly one over less of bowling per innings-6 balls.

### b) Individual quality and terminal declines

In the absence of clear senescence effects in the key performance criteria, model comparisons instead highlighted strong age-independent contributions to performance in the form of terminal declines in the last year of a career and a clear effect of longevity – the ‘age at last match’. In general, the performance of Test match bowlers deteriorated in the last year of their career, as reflected in the increase in economy rate (LMMs, fast bowler estimate = 0.126 ± 0.036, χ^2^_1_ = 12.11, p < 0.001; slow bowler estimate = 0.121 ± 0.052, χ^2^_1_ = 5.32, p = 0.021) and the decline in wicket-taking ability (GLMMs, fast bowler estimate on log-link scale = −0.110 ± 0.025, z-value = −4.32, p = < 0.001; slow bowler estimate on log-link scale = −0.188 ± 0.041, z-value = − 4.60, p < 0.001). The ‘age at last match’ covariate provided a signature of individual quality in most models (Figure 1), achieving significance in 5 of 6 cases (Supplementary Tables). The only exception to this pattern was in the wicket-taking ability of slow bowlers (GLMM, estimate on logit-link scale = 0.036 ± 0.025, z-value = 1.42, p = 0.157). Where significant, the longevity term indicated that individuals with longer careers were better than the average Test match bowler across their career, or put differently, lower quality bowlers were more likely to leave Test match cricket at a younger age.

The subsequent addition of an interaction between terminal decline term and the age at last appearance to each of the best fitting models indicated that magnitude of this terminal decline in performance was independent of the quality of bowlers, being non-significant in all cases (fast bowlers economy estimate (LMM) = 0.024 ± 0.031, χ^2^_1_ = 0.57, p = 0.45, slow bowlers economy estimate (LMM) = −0.023 ± 0.048, χ^2^_1_ = 0.22, p = 0.64; fast bowler wicket-taking estimate (GLMM) = 0.008 ± 0.022, z-value = 0.34, p = 0.73; slow bowler wicket-taking estimate (GLMM) = 0.001 ± 0.032, z-value = 0.02, p = 0.99).

### c) Multivariate bowlers

Multivariate models identified a significant, negative among-individual correlation between economy rate and wicket-taking ability in both fast and slow bowlers in models where only group-level effects were present (fast bowlers estimate = −0.27, 95% credible interval = − 0.45 – −0.08; slow bowlers estimate = −0.45, 95% credible interval = −0.64 – −0.22). After controlling for various population-level effects, both posterior correlations were reduced, but while there was little support for any correlation in fast bowlers thereafter (0.06, 95% credible interval = −0.18 –0.29), in slow bowlers there remained a significant negative correlation (−0.29, 95% credible interval = −0.55 – 0.00), suggesting that in general, the most economical slow bowlers in the modern era of Test match cricket have also been the most productive at taking wickets (see Table S4 for full model outputs). The incorporation of players that are still playing Test match cricket – or those that retired only very recently – produced very similar posterior correlations (fast bowlers estimate = 0.04, 95% credible interval = −0.17 – 0.25, slow bowlers estimate = −0.30, 95% credible interval = −0.54 – 0.05) and identified a number of players in the modern game who on their current trajectory could be considered amongst the best bowlers of the modern era (Figure S1).

## DISCUSSION

Elite players in many sporting disciplines experience declines in performance at around thirty years of age, but this has not been the case for bowlers playing Test match cricket in the modern era. Irrespective of the style of the bowler – whether fast or slow – I find that neither economy rate nor wicket-taking ability demonstrated significant age-related declines. On the contrary, while the economy rate and wicket-taking ability of slow bowlers was maintained across their career, for fast bowlers both performance metrics improved with increasing age. Despite this, a long career is not reserved for all individuals. Test cricket, as its name implies, is a testing arena in which to perform, and in most aspects of performance we find that it is only the highest quality bowlers that continue to play into their mid-to-late thirties before being replaced. Multivariate models also highlighted a clear separation between bowling styles on the basis of the relationship between economy rate and wicket-taking ability. There was no correlation among fast bowlers between the two performance metrics having controlled for various confounding effects, whereas in slow bowlers – the majority of whom are spinners – individuals with lower economy rates also tended to display higher wicket-taking ability. This finding lends credence to oft-spoken adage that ‘pressure brings wickets’ and supports the notion of a general axis of quality in international spin bowlers.

Clearly, Test match bowlers are not immune to the physiological downturns that affect elite level sportsmen and women in other dynamic sports. In part, the ability to detect senescence in performance is likely to be obscured by the strong cohort effect that manifests in only the highest quality bowlers being retained in the older age classes. Specifically, late-playing fast bowlers have above-average wicket-taking ability and bowl proportionally more overs compared to players that leave Test cricket early, while late-playing slow bowlers have above-average economy rates and likewise bowl proportionally more overs. Such cohort effects are largely unappreciated in longitudinal studies of sporting performance but are likely to be widespread, and where present, they have the potential to hinder the accurate quantification of age-related changes occurring within individuals (van de Pol and Verhulst, 2006; van de Pol and Wright, 2009). Furthermore, it is clear that any downturns in performance, whether senescent or otherwise, are highly detrimental to a cricketer’s ongoing international career. Across all but one model, bowlers were seen to experience a significant terminal decline in the last year of their Test career, at which point they were presumably dropped from the team and permanently replaced. The magnitude of the terminal declines were unaffected by the age at which a bowler finished playing, though it is possible that terminal declines are underestimated at older ages because a sizeable fraction of international cricketers retire, and will often do so ‘on a high’. As a result, players disappearing from international cricket for dips in form beyond thirty years of age were not differentiated from players who retired, which is likely to have reduced the effect size of terminal declines in elder players. It would be interesting to know whether similar cohort effects and terminal declines are prevalent in domestic cricket where selection criteria are likely to be more forgiving and where individuals are likely to be retained in teams for longer in spite of declining performance. This might allow one to pick up clearer age-related declines in performance on the domestic circuit, but on the international scene, dips in form are heavily penalised and careers are curtailed quickly thereafter.

The absence of senescence in overall performance does not preclude age-related declines in other physiological attributes. In particular, it might be expected that the speed at which fast bowlers deliver the ball declines with age. Serving speed peaks in professional tennis players at around 28 years of age (Sutter et al., 2018) and the strikeout rate of pitchers in baseball – which is thought to lean heavily on pitching speed – reaches its peak at just 23.5 years of age (Bradbury, 2009). Both of these activities bear biomechanical similarities to fast bowling which places heavy demands on joints and on arm tendons and muscles (Orchard et al., 2015). Whether or not fast bowlers do indeed bowl more slowly could not be discerned, for available speed data is not publicly available, but it should exist in other databases (e.g. CricViz) and it remains an intriguing possibility that as fast bowlers age they might compensate for declines in speed by improving their accuracy or modifying other aspects of their bowling. The reductions in economy rate with increasing age picked up by the models in this study could be indicative of improved accuracy in fast bowlers as their bowling speed declines with age.

Why is it then that various aspects of bowling performance do not senesce but rather improve with age? Bowling generally requires a high degree of motor control and so it seems likely that domain-specific expertise, deliberate practice, and increased experience (in this case of Test match cricket) are crucial to the ongoing success of bowlers. Such factors are frequently invoked to explain maintenance or improvement of high-level sporting abilities more generally (Helsen et al., 1998; Baker and Young, 2014; Careau and Wilson, 2017), and further information on training regimes and more nuanced data on delivery trajectories under Test conditions could shed light on each of these possibilities. Even so, distinguishing between the various alternatives is likely to be difficult since practice, expertise, and experience are likely to be highly inter-related. Nevertheless, a number of domain-specific characteristics that are not necessarily reliant on physical ability could be hypothesized as being important for the continued improvement of bowlers with increasing age: individuals could improve their ability to ‘work out’ individual batters and target their weaknesses, could better adapt to the highly variable conditions they experience across the world, or could develop a greater variety of deliveries that improve their likelihood of taking wickets. Another possibility is that bowlers become more adept at bowling to left-handed batters later in their career, or vice versa. Previously, Brooks et al. (2004) analysed batting performance in the 2003 Cricket World Cup and demonstrated that left-handed batters enjoyed a strategic advantage over their right handed counterparts, perhaps because of their relative rarity on the domestic circuit, with the implication that bowlers were less experienced at bowling to left handers on the international stage and thus suffered poorer returns. To my knowledge, whether such a left-handed advantage is present in the longer format of cricket remains untested, but if present, it is not inconceivable that bowlers could offset this advantage with increased practice and training as their career progresses.

The application of a multivariate modelling framework revealed an important distinction between fast and slow bowlers in the relationship between economy rate and wicket-taking ability. While fast bowlers showed no apparent relationship between the two performance metrics once other covariates are controlled for, the negative correlation in slow bowlers is suggestive of a general axis of quality whereby those slow bowlers who are more economical also tend to take more wickets. The reasons for this are intuitive-low economy rate is often thought to reflect accuracy to some degree, such that slow bowlers with lower economy rate are more accurate and thus ultimately derive greater success in the form of wickets. Conversely, the lack of equivalent trend in fast bowlers might be partly explained by an indirect trade-off between speed and accuracy at the higher velocities with which fast bowlers deliver the ball. Although any such assessment in Test cricketers would require the explicit incorporation of speed information into the multivariate framework, such a trade-off has been observed in tennis and football (van den Tillaar and Fuglstad, 2017; Sutter et al., 2018; see also Fitts, 1954) and it may have contributed towards the lack of negative association between economy-rate and wicket-taking ability here (recall that low economy rate is favourable such that any significant trade-off would be noted by a positive correlation). It is notable for example that some of the very best bowlers of the modern era in terms of wicket-taking ability are some of the very fastest of all fast bowlers (Figure 3, Figure S1), and that these individuals are rarely very economical. Even so, economy rate is not directly substitutable for accuracy, and even if it were, accuracy or skill is hard to quantify in a game where variety in terms of the trajectory and speed at which a ball is bowled can also bring rewards. As a result, testing for a correlation between speed and economy rate at broad scales – or accuracy by some measure (Feros et al., 2018) – is likely to remain elusive under match conditions.

**Figure 3.**
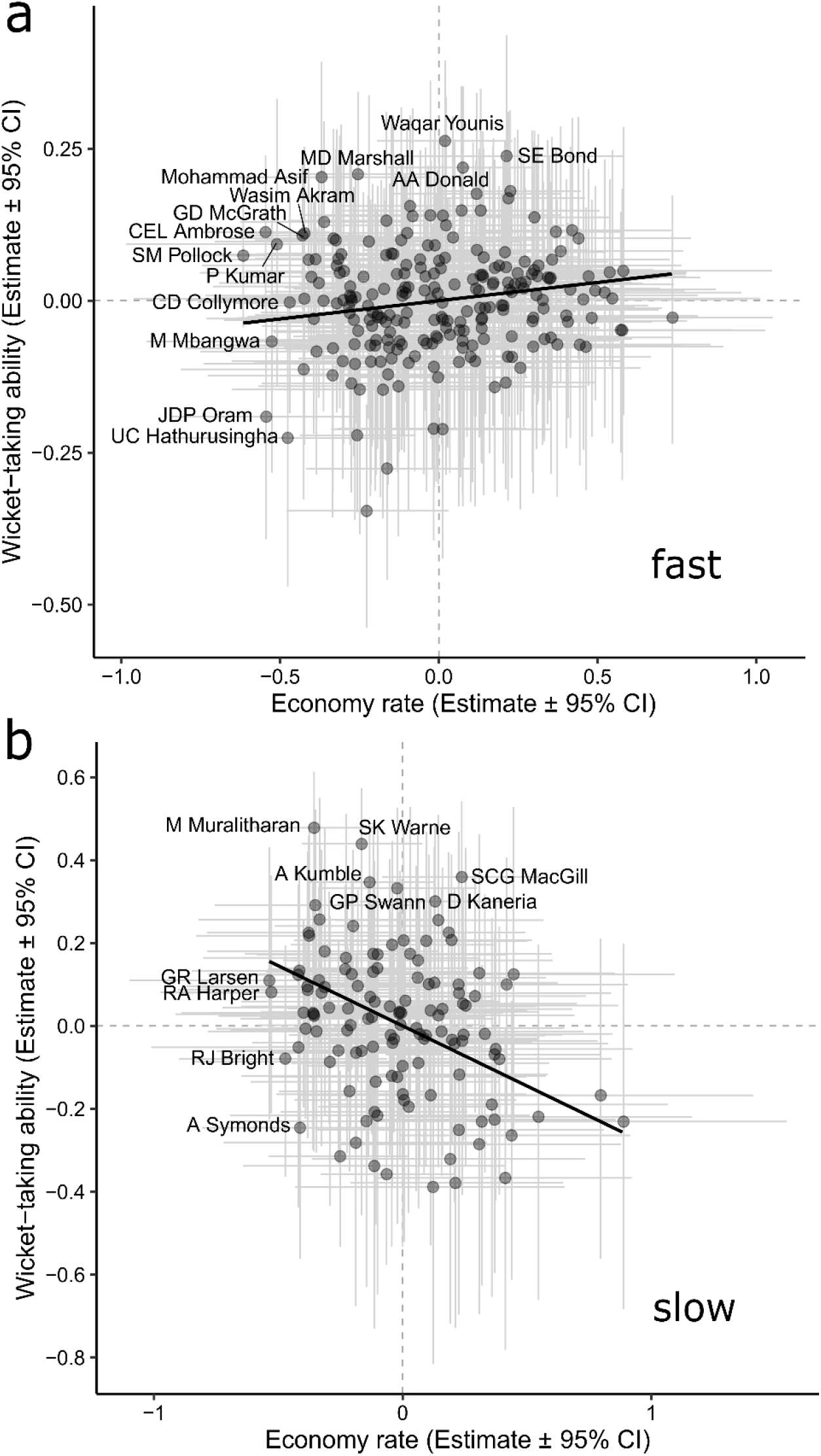
Posterior correlation between economy rate and wicket-taking ability in a) fast and b) slow bowlers, as estimated from multivariate models. Points denote the player-specific deviations in intercept for each performance metric with 95% CI (grey lines). The posterior among-individual correlation is displayed by the solid line. Estimates are taken from models that controlled for other confounding population-level effects. Individual players with particularly low economy rate or particularly high wicket-taking ability have been highlighted. For aesthetic reasons Mohammed Ashraful is not displayed in figure 3b as he represented an extreme outlier among slow bowlers.

No consideration of individual performance in cricket is complete without a discussion of rankings. Such discussions are often complicated by the need to draw comparisons across different eras (Boys and Philipson, 2019), however, the multivariate methods used in this study provide one such means of ranking Test match bowlers after having statistically controlled for temporal changes in the quality of the team and the opposition, alongside other confounding variables. If anything, the inclusion of a group-level effect of the Test playing nation in interaction with the decade will slightly devalue players from the best-performing teams through time. Nor do the models weight players according to their career longevity, which many would perhaps like to be included in any consideration of overall ranking, but these caveats noted, the approach provides a standardised means of ranking bowlers across the generations of the modern era on the basis of their individual-level intercept (their deviation from the population mean in the multivariate models; the full list of modern-era bowlers ranked by their individual-level intercepts is provided in the supplementary tables 6-9). If we consider the slow bowlers first, then known high performers are recovered at the top of the rankings: Muttiah Muralitharan (Sri Lanka) and Shane Warne (Australia) sit first and second, respectively, in terms of their wicket-taking ability, though other players retain high wicket-taking ability while also conceding very few runs per over (the upper left of figure 3b). For fast bowlers, the best bowler in terms of their wicket-taking ability is Waqar Younis (Pakistan), followed by Shane Bond (New Zealand) and Alan Donald (South Africa), though with a less clear relationship between economy rate and wicket-taking ability in fast bowlers, it could also be argued that other more economical bowlers have been equally effective operators on the international stage. The inclusion of players that are still playing Test match cricket or those that only recently stopped playing identified a number of additional top performers who on current or recent merit would be placed amongst the very best of the modern era (Fig S1). After their inclusion, for example, recently retired Dale Steyn (South Africa) becomes the best-ranked fast bowler on the basis of wicket-taking ability, being closely followed by Kagiso Rabada (South Africa), who at 24 years of age at the time of writing, is still likely to play test match cricket for several years to come.

Overall, the results of this study reiterate the need for analyses of human sporting performance to fully consider the variation occurring within and between individuals if they are to accurately capture the shape and magnitude of age trajectories. Ecological and evolutionary studies of animal populations provide a rich evidence-base to aid in this pursuit, which if employed in a human setting (Careau and Wilson, 2017), have the potential to enrich our understanding of changes in functional capacity across the human lifespan.

## ACKNOWLEDGEMENTS

JT would like to thank Leejiah Dorward, and current and former members of the University of Cambridge Zoology Department cricket team for comments that improved the manuscript.

## DATA ACCESSIBILITY

There is no funding to declare for this manuscript

## SUPPORTING INFORMATION

### Supporting Information 1 (PDF): Supplementary tables and figures

**Figure S1:** Posterior correlation between economy rate and wicket-taking ability in a) fast and b) slow bowlers, as estimated from multivariate models including Test cricketers that are still playing, or have only recently stopped playing. **Table S1:** Best fitting linear mixed effects model for the economy rate of fast bowlers and slow bowlers. **Table S2:** Best fitting generalised linear mixed effects model for the wicket-taking ability of fast bowlers and slow bowlers. **Table S3:** Best fitting generalised linear mixed effects model for the proportion of the overs bowled in the innings. **Table S4:** Results from Bayesian multivariate response models investigating the posterior correlation between economy rate and wicket-taking ability in fast bowlers. **Table S5:** Results from Bayesian multivariate response models investigating the posterior correlation between economy rate and wicket-taking ability in slow bowlers. **Table S6:** Rankings for the fast bowlers of modern era (post-1973) Test cricket on the basis of their wicket-taking ability. **Table S7:** Rankings for the fast bowlers of modern era (post-1973) Test cricket on the basis of their economy rate. **Table S8:** Rankings for the slow bowlers of modern era (post-1973) Test cricket on the basis of their wicket-taking ability. **Table S9:** Rankings for the slow bowlers of modern era (post-1973) Test cricket on the basis of their economy rate.

### Supporting Information 2 (XLS)

Data files used for analyses.

