## Supporting Information 1 for "Bowled over or over bowled? Age-related changes in the performance of bowlers in Test match cricket"

Jack Thorley

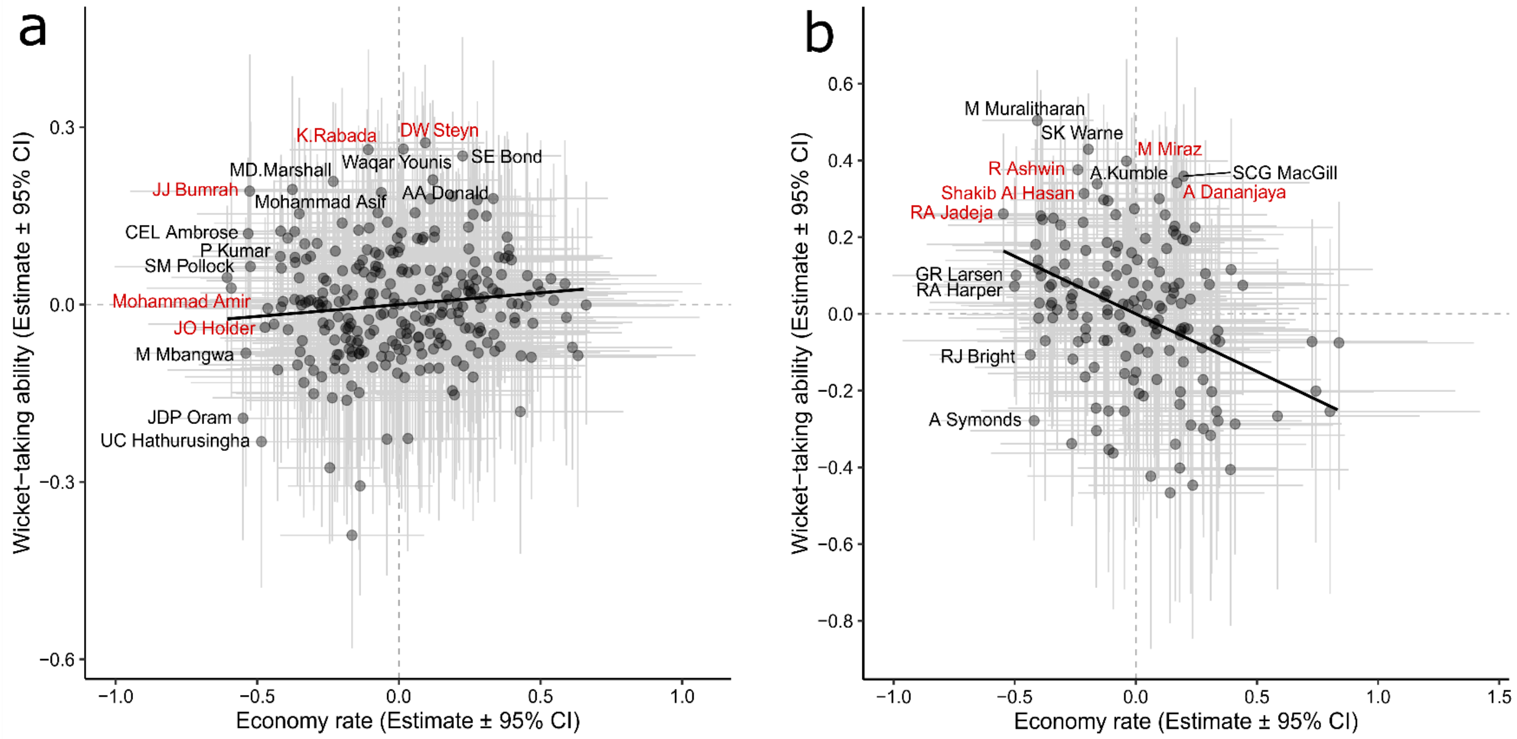

**Figure S1. Posterior correlation between economy rate and wicket-taking ability in a) fast and b) slow bowlers, as estimated from multivariate models, including Test cricketers that are still playing, or have only recently stopped playing.** Points denote the player-specific deviations in intercept for each performance metric with 95% CI (grey lines). The posterior among-individual correlation is displayed by the solid line. Estimates are taken from models that controlled for other confounding population-level effects. Individual players with particularly low economy rate or particularly high wicket-taking ability have been highlighted; current players or bowlers that have only recently retired are coloured red.

**Table S1: Best fitting linear mixed effects model for the economy rate of fast bowlers and slow bowlers.** Significance (α < 0.05) of fixed effects was assessed by likelihood ratio tests when respective terms were dropped from the main model. We do not present the significance of main effects when present in interactions. Continuous variables were z-score transformed prior to model fitting. 95% confidence intervals for the fixed effects means were calculated by parametric bootstrapping.

**FAST BOWLERS ECONOMY**

| **Fixed Effect** | **Estimate** | **SE** | **95% CI** | **t-value** | **$X^2$, df** | **p- value** |
|---|---|---|---|---|---|---|
| Intercept | 3.094 | 0.055 | 2.984 – 3.201 | | | |
| Age | -0.105 | 0.025 | -0.155 – -0.057 | -4.19 | | |
| Innings- 2 | 0.063 | 0.032 | -0.003 – 0.126 | 1.96 | | |
| 3 | 0.049 | 0.027 | 0.001 – 0.103 | 1.79 | | |
| 4 | 0.301 | 0.036 | 0.222 – 0.370 | 8.26 | | |
| Home or Away- Home | -0.120 | 0.028 | -0.177 – -0.071 | -4.28 | 18.24, 1 | < 0.001 |
| Terminal Effect | 0.126 | 0.036 | 0.055 – 0.194 | 3.50 | 12.11, 1 | < 0.001 |
| Last Game Age | -0.080 | 0.030 | -0.135 – 0.021 | -2.68 | 6.69, 1 | 0.010 |
| Age:Innings2 | 0.010 | 0.029 | -0.049 – 0.069 | 0.35 | 12.46, 3 | 0.006 |
| Age:Innings3 | -0.013 | 0.027 | -0.066 – 0.040 | -0.46 | | |
| Age:Innings4 | -0.094 | 0.033 | -0.158 – -0.026 | -2.88 | | |

| **Random Effect** | **Variance** | **Std Dev** |
|---|---|---|
| PlayerID (n = 227) | 0.104 | 0.323 |
| MatchID (n = 2581) | 0.196 | 0.443 |
| Opposition:Decade (n = 42) | 0.050 | 0.220 |
| Country:Decade (n = 40) | 0.006 | 0.077 |
| Residual | 1.113 | 1.055 |

**SLOW BOWLERS ECONOMY**

| **Fixed Effect** | **Estimate** | **SE** | **95% CI** | **t-value** | **$X^2$, df** | **p- value** |
|---|---|---|---|---|---|---|
| Intercept | 3.064 | 0.078 | 2.912 – 3.210 | | | |
| Innings- 2 | -0.178 | 0.050 | -0.273 – -0.083 | -3.59 | 31.26, 3 | < 0.001 |
| 3 | -0.067 | 0.047 | -0.162 – 0.021 | -1.43 | | |
| 4 | 0.108 | 0.057 | -0.011 – 0.220 | 1.88 | | |
| Home or Away- Home | -0.169 | 0.041 | -0.245 – -0.092 | -4.12 | 16.86, 1 | < 0.001 |
| Terminal Effect | 0.121 | 0.053 | 0.010 – 0.216 | 2.32 | 5.32, 1 | 0.021 |
| Last Game Age | -0.155 | 0.039 | -0.231 – -0.077 | -3.99 | 15.44, 1 | < 0.001 |

| **Random Effect** | **Variance** | **Std Dev** |
|---|---|---|
| PlayerID (n =128) | 0.142 | 0.388 |
| MatchID (n = 2044) | 0.151 | 0.376 |
| Opposition:Decade (n = 42) | 0.082 | 0.285 |
| Country:Decade (n = 41) | 0.024 | 0.155 |
| Residual | 1.420 | 1.191 |

**Table S2: Best fitting generalised linear mixed effects model for the wicket-taking ability of fast bowlers and slow bowlers.** Models were fitted to a negative binomial error distribution with a zero-inflation parameter applied across all data points (~1). 95% confidence intervals for the fixed effects means were calculated via the Wald method.

**FAST BOWLERS WICKETS**

| **Fixed Effect** | **Estimate** | **SE** | **95% CI** | **z-value** | **p- value** |
|---|---|---|---|---|---|
| Intercept | 0.531 | 0.029 | 0.475 – 0.587 | | |
| Age | -0.024 | 0.017 | -0.057 – 0.008 | -1.46 | 0.145 |
| Innings- 2 | -0.014 | 0.020 | -0.053 – 0.024 | -0.73 | 0.463 |
| 3 | -0.041 | 0.021 | -0.082 – 0.000 | -1.95 | 0.051 |
| 4 | -0.150 | 0.027 | -0.203 – -0.097 | -5.56 | < 0.001 |
| Home or Away- Home | 0.069 | 0.016 | 0.038 – 0.100 | 4.41 | < 0.001 |
| Overs | 0.324 | 0.009 | 0.307 – 0.341 | 36.95 | < 0.001 |
| Terminal Effect | -0.110 | 0.026 | -0.160 – 0.060 | -4.32 | < 0.001 |
| Last Game Age | 0.040 | 0.017 | 0.007 – 0.073 | 2.36 | 0.018 |
| Age:Innings2 | 0.039 | 0.020 | 0.001 – 0.078 | 2.02 | 0.043 |
| Age:Innings3 | 0.045 | 0.021 | 0.005 – 0.086 | 2.18 | 0.029 |
| Age:Innings4 | 0.082 | 0.025 | 0.032 – 0.132 | 3.23 | 0.001 |
| **Zero-inflation Intercept** | -4.97 | 0.843 | | -5.90 | < 0.001 |
| **Random Effect** | **Variance** | **Std Dev** | | | |
| PlayerID (n =227) | 0.018 | 0.134 | | | |
| Opposition:Decade (n = 42) | 0.006 | 0.078 | | | |
| Country:Decade (n = 40) | 0.006 | 0.075 | | | |

**SLOW BOWLERS WICKETS**

| **Fixed Effect** | **Estimate** | **SE** | **95% CI** | **z-value** | **p- value** |
|---|---|---|---|---|---|
| Intercept | 0.284 | 0.047 | 0.192 – 0.376 | | |
| Innings- 2 | 0.044 | 0.033 | -0.020 – 0.108 | 1.34 | 0.180 |
| 3 | 0.135 | 0.033 | 0.071 – 0.199 | 4.14 | < 0.001 |
| 4 | 0.075 | 0.041 | -0.004 – 0.154 | 1.84 | 0.066 |
| Home or Away- Home | 0.083 | 0.026 | 0.033 – 0.133 | 3.23 | 0.001 |
| Overs | 0.414 | 0.014 | 0.387 – 0.441 | 30.20 | < 0.001 |
| Terminal Effect | -0.163 | 0.036 | -0.234 – -0.092 | -4.51 | < 0.001 |
| Last Game Age | 0.036 | 0.025 | -0.014 – 0.086 | 1.42 | 0.157 |
| **Zero-inflation Intercept** | -15.60 | 1243.6 | | -0.01 | 0.99 |
| **Random Effect** | **Variance** | **Std Dev** | | | |
| PlayerID (n =128) | 0.054 | 0.233 | | | |
| Opposition:Decade (n = 41) | 0.022 | 0.148 | | | |
| Country:Decade (n = 40) | 0.006 | 0.080 | | | |

**Table S3: Best fitting generalised linear mixed effects model for the proportion of the overs bowled in the innings.** Models were fitted to a binomial error distribution with the number of overs bowler by the bowler as the numerator, and the total number of overs in the innings set as the denominator. 95% confidence intervals for the fixed effects means were calculated by parametric bootstrapping.

**FAST BOWLERS OVERS**

| **Fixed Effect** | **Estimate** | **SE** | **95% CI** | **z-value** | **p- value** |
|---|---|---|---|---|---|
| Intercept | -1.578 | 0.017 | -1.615 – -1.545 | | |
| Age | 0.011 | 0.005 | -0.003 – 0.004 | 1.54 | 0.124 |
| $Age^2$ | -0.032 | 0.003 | -0.004 – -0.022 | -6.48 | < 0.001 |
| Innings- 2 | -0.020 | 0.008 | -0.037 – -0.007 | -2.35 | 0.019 |
| 3 | -0.087 | 0.007 | -0.099 – -0.073 | -12.61 | < 0.001 |
| 4 | -0.050 | 0.011 | -0.070 – -0.031 | -4.64 | < 0.001 |
| Home or Away- Home | -0.027 | 0.008 | -0.037 – -0.007 | -2.85 | 0.004 |
| Terminal Effect | -0.049 | 0.011 | -0.049 – -0.005 | 2.41 | 0.016 |
| Last Game Age | 0.078 | 0.014 | 0.052 – 0.106 | 5.54 | < 0.001 |
| Age: Last Game Age | 0.034 | 0.007 | 0.019 – 0.048 | 4.62 | < 0.001 |
| $Age^2$: Last Game Age | 0.003 | 0.003 | -0.002 – 0.008 | 1.24 | 0.215 |

| **Random Effect** | **Variance** | **Std Dev** |
|---|---|---|
| MatchID (n = 2568) | 0.020 | 0.144 |
| PlayerID (n =219) | 0.023 | 0.152 |
| Opposition:Decade (n = 42) | 0.000 | 0.018 |
| Country:Decade (n = 40) | 0.004 | 0.066 |

**SLOW BOWLERS OVERS**

| **Fixed Effect** | **Estimate** | **SE** | **95% CI** | **z-value** | **p- value** |
|---|---|---|---|---|---|
| Intercept | -1.725 | 0.046 | -1.812 – -1.634 | | |
| Age | -0.035 | 0.018 | -0.072 – 0.003 | -1.92 | 0.055 |
| $Age^2$ | -0.038 | 0.012 | -0.006 – -0.015 | -3.16 | 0.002 |
| Innings- 2 | 0.092 | 0.016 | 0.059 – 0.126 | 5.75 | < 0.001 |
| 3 | 0.192 | 0.012 | 0.167 – 0.216 | 15.43 | < 0.001 |
| 4 | 0.250 | 0.020 | 0.205 – 0.293 | 12.37 | < 0.001 |
| Home or Away- Home | -0.015 | 0.013 | -0.040 – 0.012 | -1.10 | 0.273 |
| Terminal Effect | -0.016 | 0.021 | -0.056 – -0.022 | -0.78 | 0.438 |
| Last Game Age | 0.149 | 0.037 | 0.080 – 0.226 | 4.06 | < 0.001 |
| Age:Innings2 | 0.023 | 0.011 | 0.002 – 0.049 | 2.09 | 0.037 |
| Age:Innings3 | 0.043 | 0.010 | 0.023 – 0.065 | 4.27 | < 0.001 |
| Age:Innings4 | 0.021 | 0.015 | 0.010 – 0.051 | 1.47 | 0.141 |
| $Age^2$:Innings2 | -0.013 | 0.009 | -0.030 – 0.004 | -1.49 | 0.136 |
| $Age^2$:Innings3 | -0.007 | 0.008 | -0.022 – 0.010 | -0.86 | 0.392 |
| $Age^2$:Innings4 | -0.016 | 0.012 | -0.041 – 0.008 | -1.36 | 0.173 |
| Age: Last Game Age | 0.085 | 0.017 | 0.051 – 0.121 | 4.93 | < 0.001 |
| $Age^2$: Last Game Age | -0.007 | 0.006 | -0.020 – 0.007 | -1.12 | 0.264 |

| **Random Effect** | **Variance** | **Std Dev** |
|---|---|---|
| MatchID (n = 2044) | 0.053 | 0.230 |
| PlayerID (n =128) | 0.145 | 0.381 |
| Opposition:Decade (n =42) | 0.001 | 0.035 |
| Country:Decade (n =41) | 0.018 | 0.135 |

**Table S4: Results from Bayesian multivariate response models investigating the posterior correlation between economy rate and wicket-taking ability in fast bowlers.** Economy rate was fitted to a Gaussian distribution, wicket-taking ability to a negative binomial distribution. Estimates for the group-level and the population-level effects provide the standard deviation and the mean, respectively. Estimates for the wicket-taking ability model are provided on the link (log) scale.

| Term | Estimate | Lower 95% CI | Upper 95% CI |
|---|---|---|---|
| ***Group-level effects*** | | | |
| **PlayerID** | | | |
| Economy rate: Intercept | 0.33 | 0.28 | 0.38 |
| Wicket-taking ability: Intercept | 0.14 | 0.11 | 0.16 |
| Cor(Economy-Wicket-taking) | 0.06 | -0.18 | 0.29 |
| **Match ID** | | | |
| Economy rate: Intercept | 0.44 | 0.41 | 0.47 |
| **Opposition:decade** | | | |
| Economy rate: Intercept | 0.24 | 0.17 | 0.32 |
| Wicket-taking ability: Intercept | 0.08 | 0.06 | 0.11 |
| **Country:decade** | | | |
| Economy rate: Intercept | 0.11 | 0.03 | 0.20 |
| Wicket-taking ability: Intercept | 0.10 | 0.05 | 0.14 |
| ***Population-level effects*** | | | |
| Economy rate: Intercept | 3.12 | 3.01 | 3.23 |
| Economy rate: Age | -0.13 | -0.17 | -0.08 |
| Economy rate: Home or away-Home | -0.12 | -0.18 | -0.07 |
| Economy rate: Innings2 | 0.06 | 0.00 | 0.13 |
| Economy rate: Innings3 | 0.05 | -0.01 | 0.10 |
| Economy rate: Innings4 | 0.30 | 0.23 | 0.37 |
| Economy rate: Terminal effect | 0.15 | 0.09 | 0.22 |
| Economy rate: Age:Innings2 | 0.01 | -0.05 | 0.06 |
| Economy rate: Age:Innings3 | -0.01 | -0.07 | 0.04 |
| Economy rate: Age:Innings4 | -0.09 | -0.16 | -0.03 |
| Wicket-taking ability: Intercept | 0.51 | 0.45 | 0.57 |
| Wicket-taking ability: Age | -0.01 | -0.04 | 0.02 |
| Wicket-taking ability: Home or away-Home | 0.07 | 0.04 | 0.10 |
| Wicket-taking ability: Innings2 | -0.01 | -0.05 | 0.02 |
| Wicket-taking ability: Innings3 | -0.04 | -0.08 | 0.00 |
| Wicket-taking ability: Innings4 | -0.15 | -0.20 | -0.10 |
| Wicket-taking ability: Overs | 0.33 | 0.31 | 0.34 |
| Wicket-taking ability: Terminal effect | -0.13 | -0.18 | -0.08 |
| Wicket-taking ability: Age:Innings2 | 0.04 | 0.00 | 0.08 |
| Wicket-taking ability: Age:Innings3 | 0.05 | 0.01 | 0.09 |
| Wicket-taking ability: Age:Innings4 | 0.08 | 0.03 | 0.13 |
| Sigma- Economy rate | 1.06 | 1.04 | 1.07 |
| Shape parameter – Wicket-taking ability | 10.81 | 8.94 | 13.20 |

**Table S5: Results from Bayesian multivariate response models investigating the posterior correlation between economy rate and wicket-taking ability in slow bowlers.** Economy rate was fitted to a Gaussian distribution, wicket-taking ability to a negative binomial distribution. Estimates for the group-level and the population-level effects provide the standard deviation and the mean, respectively. Estimates for the wicket-taking ability model are provided on the link (log) scale.

| Term | Estimate | Lower 95% CI | Upper 95% CI |
|---|---|---|---|
| ***Group-level effects*** | | | |
| **PlayerID** | | | |
| Economy rate: Intercept | 0.41 | 0.33 | 0.50 |
| Wicket-taking ability: Intercept | 0.24 | 0.19 | 0.30 |
| Cor(Economy-Wicket-taking) | -0.29 | -0.55 | 0.00 |
| **Match ID** | | | |
| Economy rate: Intercept | 0.39 | 0.32 | 0.45 |
| **Opposition:decade** | | | |
| Economy rate: Intercept | 0.30 | 0.22 | 0.40 |
| Wicket-taking ability: Intercept | 0.16 | 0.11 | 0.22 |
| **Country:decade** | | | |
| Economy rate: Intercept | 0.15 | 0.03 | 0.28 |
| Wicket-taking ability: Intercept | 0.08 | 0.01 | 0.15 |
| ***Population-level effects*** | | | |
| Economy rate: Intercept | 3.11 | 2.96 | 3.27 |
| Economy rate: Home or away-Home | -0.16 | -0.24 | -0.09 |
| Economy rate: Innings2 | -0.18 | -0.28 | -0.08 |
| Economy rate: Innings3 | -0.07 | -0.16 | 0.03 |
| Economy rate: Innings4 | 0.11 | -0.00 | 0.22 |
| Economy rate: Terminal effect | 0.13 | 0.03 | 0.24 |
| Wicket-taking ability: Intercept | 0.27 | 0.17 | 0.36 |
| Wicket-taking ability: Home or away-Home | 0.08 | 0.03 | 0.13 |
| Wicket-taking ability: Innings2 | 0.05 | -0.02 | 0.11 |
| Wicket-taking ability: Innings3 | 0.14 | 0.07 | 0.20 |
| Wicket-taking ability: Innings4 | 0.08 | -0.00 | 0.15 |
| Wicket-taking ability: Overs | 0.41 | 0.39 | 0.44 |
| Wicket-taking ability: Terminal effect | -0.17 | -0.24 | -0.10 |
| Sigma- Economy rate | 1.19 | 1.16 | 1.22 |
| Shape parameter – Wicket-taking ability | 8.73 | 6.85 | 11.36 |

**Table S6: Rankings for the fast bowlers of modern era (post-1973) Test cricket on the basis of their wicket-taking ability.** Estimates represent the player-specific intercepts extracted from the multivariate response models that controlled for various confounding effects- including the number of overs bowled per innings. Players that were still thought to be playing Test cricket on or around June 2019 are not included.

| Rank | PlayerName | Country | Wickets | lower_95CI | upper_95CI |
|---|---|---|---|---|---|
| 1 | Waqar.Younis | Pakistan | 0.2628 | 0.1289 | 0.3947 |
| 2 | SE.Bond | New Zealand | 0.2380 | 0.0458 | 0.4369 |
| 3 | AA.Donald | South Africa | 0.2190 | 0.0832 | 0.3528 |
| 4 | MD.Marshall | West Indies | 0.2080 | 0.0756 | 0.3439 |
| 5 | Mohammad.Asif | Pakistan | 0.2033 | 0.0217 | 0.3931 |
| 6 | MG.Johnson | Australia | 0.1803 | 0.0523 | 0.3132 |
| 7 | Shoaib.Akhtar | Pakistan | 0.1758 | 0.0194 | 0.3419 |
| 8 | D.Gough | England | 0.1685 | 0.0240 | 0.3110 |
| 9 | M.Morkel | South Africa | 0.1557 | 0.0134 | 0.3049 |
| 10 | DW.Headley | England | 0.1484 | -0.0549 | 0.3528 |
| 11 | JL.Pattinson | Australia | 0.1483 | -0.0543 | 0.3487 |
| 12 | DE.Bollinger | Australia | 0.1405 | -0.0733 | 0.3622 |
| 13 | M.Ntini | South Africa | 0.1400 | 0.0153 | 0.2684 |
| 14 | GR.Dilley | England | 0.1391 | -0.0344 | 0.3161 |
| 15 | CL.Cairns | New Zealand | 0.1372 | -0.0021 | 0.2794 |
| 16 | CEH.Croft | West Indies | 0.1317 | -0.0441 | 0.3108 |
| 17 | IR.Bishop | West Indies | 0.1295 | -0.0295 | 0.2954 |
| 18 | JJC.Lawson | West Indies | 0.1241 | -0.0837 | 0.3356 |
| 19 | SL.Malinga | SriLanka | 0.1159 | -0.0684 | 0.3011 |
| 20 | B.Lee | Australia | 0.1134 | -0.0174 | 0.2503 |
| 21 | CEL.Ambrose | West Indies | 0.1128 | -0.0125 | 0.2395 |
| 22 | RJ.Harris | Australia | 0.1113 | -0.0662 | 0.2868 |
| 23 | GD.McGrath | Australia | 0.1110 | 0.0004 | 0.2197 |
| 24 | Wasim.Akram | Pakistan | 0.1075 | -0.0155 | 0.2339 |
| 25 | LS.Pascoe | Australia | 0.1040 | -0.0931 | 0.3032 |
| 26 | BA.Reid | Australia | 0.1025 | -0.0760 | 0.2830 |
| 27 | KTGD.Prasad | SriLanka | 0.1025 | -0.0962 | 0.3028 |
| 28 | BN.Schultz | South Africa | 0.0996 | -0.1211 | 0.3183 |
| 29 | J.Garner | West Indies | 0.0995 | -0.0458 | 0.2494 |
| 30 | AH.Gray | West Indies | 0.0974 | -0.1376 | 0.3381 |
| 31 | P.Kumar | India | 0.0928 | -0.1421 | 0.3323 |
| 32 | SP.Jones | England | 0.0911 | -0.1056 | 0.2988 |
| 33 | CJ.McDermott | Australia | 0.0909 | -0.0419 | 0.2222 |
| 34 | M.Hayward | South Africa | 0.0835 | -0.1242 | 0.2922 |
| 35 | ALF.de.Mel | SriLanka | 0.0814 | -0.1338 | 0.2873 |
| 36 | ST.Finn | England | 0.0798 | -0.1060 | 0.2632 |
| 37 | CT.Tremlett | England | 0.0777 | -0.1302 | 0.2942 |
| 38 | WW.Daniel | West Indies | 0.0769 | -0.1416 | 0.3041 |
| 39 | Shabbir.Ahmed | Pakistan | 0.0764 | -0.1346 | 0.2903 |
| 40 | PS.de.Villiers | South Africa | 0.0749 | -0.1171 | 0.2735 |

| | | | | | |
|---|---|---|---|---|---|
| 41 | SM.Pollock | South Africa | 0.0742 | -0.0440 | 0.1941 |
| 42 | S.Sreesanth | India | 0.0713 | -0.1172 | 0.2507 |
| 43 | AR.Caddick | England | 0.0706 | -0.0647 | 0.2073 |
| 44 | ARC.Fraser | England | 0.0689 | -0.0849 | 0.2217 |
| 45 | CA.Walsh | West Indies | 0.0682 | -0.0433 | 0.1788 |
| 46 | IK.Pathan | India | 0.0680 | -0.1151 | 0.2478 |
| 47 | AM.Blignaut | Zimbabwe | 0.0676 | -0.1393 | 0.2696 |
| 48 | HH.Streak | Zimbabwe | 0.0673 | -0.0838 | 0.2189 |
| 49 | Azeem.Hafeez | Pakistan | 0.0659 | -0.1322 | 0.2705 |
| 50 | DK.Morrison | New Zealand | 0.0642 | -0.0970 | 0.2204 |
| 51 | J.Srinath | India | 0.0588 | -0.0825 | 0.1985 |
| 52 | KJ.Abbott | South Africa | 0.0577 | -0.1612 | 0.2821 |
| 53 | Z.Khan | India | 0.0577 | -0.0722 | 0.1885 |
| 54 | G.Onions | England | 0.0577 | -0.1660 | 0.2741 |
| 55 | DAJ.Bracewell | New Zealand | 0.0552 | -0.1452 | 0.2504 |
| 56 | RJ.Ratnayake | SriLanka | 0.0545 | -0.1305 | 0.2479 |
| 57 | KCG.Benjamin | West Indies | 0.0539 | -0.1259 | 0.2322 |
| 58 | MG.Hughes | Australia | 0.0533 | -0.0879 | 0.1968 |
| 59 | Wahab.Riaz | Pakistan | 0.0503 | -0.1546 | 0.2610 |
| 60 | CEL.Stuart | West Indies | 0.0490 | -0.1880 | 0.2862 |
| 61 | SR.Clark | Australia | 0.0486 | -0.1317 | 0.2280 |
| 62 | FH.Edwards | West Indies | 0.0466 | -0.1076 | 0.1995 |
| 63 | T.Panyangara | Zimbabwe | 0.0459 | -0.1774 | 0.2716 |
| 64 | Mohsin.Kamal | Pakistan | 0.0454 | -0.1878 | 0.2818 |
| 65 | GF.Lawson | Australia | 0.0451 | -0.1124 | 0.2019 |
| 66 | TM.Alderman | Australia | 0.0447 | -0.1097 | 0.2025 |
| 67 | M.Zondeki | South Africa | 0.0438 | -0.1918 | 0.2805 |
| 68 | SB.Doull | New Zealand | 0.0438 | -0.1339 | 0.2194 |
| 69 | MJ.Hoggard | England | 0.0427 | -0.0969 | 0.1817 |
| 70 | IE.O'Brien | New Zealand | 0.0422 | -0.1504 | 0.2380 |
| 71 | NG.Cowans | England | 0.0405 | -0.1710 | 0.2518 |
| 72 | BP.Patterson | West Indies | 0.0400 | -0.1406 | 0.2173 |
| 73 | RM.Ellison | England | 0.0400 | -0.1815 | 0.2523 |
| 74 | JN.Gillespie | Australia | 0.0395 | -0.0958 | 0.1789 |
| 75 | Aamer.Nazir | Pakistan | 0.0393 | -0.1965 | 0.2783 |
| 76 | WPUJC.Vaas | SriLanka | 0.0392 | -0.0905 | 0.1713 |
| 77 | Sohail.Khan | Pakistan | 0.0383 | -0.2029 | 0.2645 |
| 78 | JE.Taylor | West Indies | 0.0377 | -0.1299 | 0.1952 |
| 79 | C.Sharma | India | 0.0362 | -0.1639 | 0.2385 |
| 80 | AJ.Tudor | England | 0.0351 | -0.1898 | 0.2604 |
| 81 | DG.Cork | England | 0.0311 | -0.1248 | 0.1854 |
| 82 | CS.Martin | New Zealand | 0.0297 | -0.1173 | 0.1735 |
| 83 | L.Balaji | India | 0.0297 | -0.2034 | 0.2677 |
| 84 | BW.Hilfenhaus | Australia | 0.0291 | -0.1594 | 0.2113 |
| 85 | RP.Singh | India | 0.0285 | -0.1821 | 0.2404 |
| 86 | DV.Lawrence | England | 0.0270 | -0.2111 | 0.2688 |
| 87 | IG.Butler | New Zealand | 0.0269 | -0.2062 | 0.2540 |

| | | | | | |
|---|---|---|---|---|---|
| 88 | CK.Langeveldt | South Africa | 0.0268 | -0.2097 | 0.2652 |
| 89 | ST.Clarke | West Indies | 0.0234 | -0.1934 | 0.2419 |
| 90 | PM.Siddle | Australia | 0.0220 | -0.1288 | 0.1692 |
| 91 | BT.Watambwa | Zimbabwe | 0.0219 | -0.2232 | 0.2619 |
| 92 | DW.Fleming | Australia | 0.0218 | -0.1672 | 0.2073 |
| 93 | T.Thushara | SriLanka | 0.0203 | -0.2077 | 0.2336 |
| 94 | Junaid.Khan | Pakistan | 0.0198 | -0.1913 | 0.2310 |
| 95 | A.Nel | South Africa | 0.0194 | -0.1441 | 0.1852 |
| 96 | SI.Mahmood | England | 0.0183 | -0.2254 | 0.2538 |
| 97 | KR.Pushpakumara | SriLanka | 0.0178 | -0.1902 | 0.2210 |
| 98 | M.Dillon | West Indies | 0.0169 | -0.1481 | 0.1781 |
| 99 | CRD.Fernando | SriLanka | 0.0166 | -0.1638 | 0.1884 |
| 100 | N.Phillip | West Indies | 0.0163 | -0.2252 | 0.2500 |
| 101 | A.Kuruvilla | India | 0.0149 | -0.2220 | 0.2406 |
| 102 | Umar.Gul | Pakistan | 0.0142 | -0.1475 | 0.1792 |
| 103 | DH.Brain | Zimbabwe | 0.0135 | -0.2091 | 0.2325 |
| 104 | SJ.Harmison | England | 0.0122 | -0.1311 | 0.1532 |
| 105 | FA.Rose | West Indies | 0.0119 | -0.1878 | 0.2149 |
| 106 | DR.Tuffey | New Zealand | 0.0105 | -0.1763 | 0.1922 |
| 107 | Talha.Jubair | Bangladesh | 0.0091 | -0.2437 | 0.2546 |
| 108 | Mashrafe.Mortaza | Bangladesh | 0.0083 | -0.1909 | 0.1986 |
| 109 | DT.Hondo | Zimbabwe | 0.0082 | -0.2376 | 0.2455 |
| 110 | GC.Small | England | 0.0058 | -0.2019 | 0.2109 |
| 111 | AIC.Dodemaide | Australia | 0.0051 | -0.2156 | 0.2227 |
| 112 | VR.Aaron | India | 0.0037 | -0.2349 | 0.2482 |
| 113 | JR.Hammond | Australia | 0.0033 | -0.2375 | 0.2533 |
| 114 | DJ.Terbrugge | South Africa | 0.0031 | -0.2274 | 0.2300 |
| 115 | DA.Stirling | New Zealand | 0.0021 | -0.2456 | 0.2460 |
| 116 | Robiul.Islam | Bangladesh | 0.0011 | -0.2377 | 0.2328 |
| 117 | MA.Ealham | England | 0.0009 | -0.2352 | 0.2436 |
| 118 | PAJ.DeFreitas | England | 0.0000 | -0.1580 | 0.1575 |
| 119 | DJ.Nash | New Zealand | 0.0000 | -0.1826 | 0.1756 |
| 120 | CD.Collymore | West Indies | -0.0028 | -0.1849 | 0.1708 |
| 121 | GF.Labrooy | SriLanka | -0.0030 | -0.2323 | 0.2242 |
| 122 | NA.Foster | England | -0.0043 | -0.1833 | 0.1724 |
| 123 | JEC.Franklin | New Zealand | -0.0075 | -0.1869 | 0.1756 |
| 124 | MB.Owens | New Zealand | -0.0082 | -0.2420 | 0.2215 |
| 125 | RJ.Sidebottom | England | -0.0083 | -0.1943 | 0.1738 |
| 126 | N.Kapil.Dev | India | -0.0094 | -0.1302 | 0.1123 |
| 127 | PDRL.Perera | SriLanka | -0.0109 | -0.2509 | 0.2212 |
| 128 | RMS.Eranga | SriLanka | -0.0118 | -0.2243 | 0.1952 |
| 129 | PT.Collins | West Indies | -0.0121 | -0.1861 | 0.1574 |
| 130 | UWMBCA.Welegedara | SriLanka | -0.0127 | -0.2238 | 0.1959 |
| 131 | MI.Black | West Indies | -0.0151 | -0.2684 | 0.2300 |
| 132 | RD.King | West Indies | -0.0174 | -0.2246 | 0.1816 |
| 133 | R.Rampaul | West Indies | -0.0190 | -0.2299 | 0.1831 |
| 134 | CR.Miller | Australia | -0.0200 | -0.2103 | 0.1720 |

| | | | | | |
|---|---|---|---|---|---|
| 135 | Harbhajan.Singh | India | -0.0201 | -0.1413 | 0.1040 |
| 136 | JR.Ratnayeke | SriLanka | -0.0230 | -0.2281 | 0.1878 |
| 137 | AtaNAurNARehman | Pakistan | -0.0237 | -0.2426 | 0.1934 |
| 138 | A.Sanford | West Indies | -0.0245 | -0.2428 | 0.1998 |
| 139 | CG.Rackemann | Australia | -0.0268 | -0.2443 | 0.1884 |
| 140 | WW.Davis | West Indies | -0.0269 | -0.2375 | 0.1834 |
| 141 | Shahadat.Hossain | Bangladesh | -0.0279 | -0.2353 | 0.1737 |
| 142 | LE.Plunkett | England | -0.0279 | -0.2412 | 0.1747 |
| 143 | HK.Olonga | Zimbabwe | -0.0284 | -0.2239 | 0.1664 |
| 144 | CJ.Jordan | England | -0.0285 | -0.2691 | 0.1988 |
| 145 | TJ.Friend | Zimbabwe | -0.0286 | -0.2685 | 0.2035 |
| 146 | WKM.Benjamin | West Indies | -0.0298 | -0.2189 | 0.1619 |
| 147 | MR.Whitney | Australia | -0.0310 | -0.2465 | 0.1746 |
| 148 | TCB.Fernando | SriLanka | -0.0311 | -0.2648 | 0.2053 |
| 149 | DNT.Zoysa | SriLanka | -0.0314 | -0.2222 | 0.1557 |
| 150 | CE.Cuffy | West Indies | -0.0315 | -0.2560 | 0.1868 |
| 151 | Saleem.Jaffar | Pakistan | -0.0318 | -0.2475 | 0.1718 |
| 152 | BKV.Prasad | India | -0.0323 | -0.2106 | 0.1441 |
| 153 | NAM.McLean | West Indies | -0.0333 | -0.2411 | 0.1675 |
| 154 | CEW.Silverwood | England | -0.0337 | -0.2824 | 0.2080 |
| 155 | DE.Malcolm | England | -0.0347 | -0.1931 | 0.1283 |
| 156 | SB.O'Connor | New Zealand | -0.0349 | -0.2301 | 0.1646 |
| 157 | VRV.Singh | India | -0.0349 | -0.2902 | 0.2131 |
| 158 | VC.Drakes | West Indies | -0.0359 | -0.2492 | 0.1697 |
| 159 | TT.Bresnan | England | -0.0414 | -0.2396 | 0.1569 |
| 160 | DK.Liyanage | SriLanka | -0.0445 | -0.2862 | 0.1967 |
| 161 | CPH.Ramanayake | SriLanka | -0.0449 | -0.2560 | 0.1544 |
| 162 | AJ.Bichel | Australia | -0.0482 | -0.2495 | 0.1464 |
| 163 | KD.Mills | New Zealand | -0.0484 | -0.2506 | 0.1462 |
| 164 | JDS.Neesham | New Zealand | -0.0487 | -0.2954 | 0.1847 |
| 165 | Shahid.Nazir | Pakistan | -0.0494 | -0.2578 | 0.1561 |
| 166 | PR.Reiffel | Australia | -0.0496 | -0.2205 | 0.1206 |
| 167 | A.Nehra | India | -0.0517 | -0.2633 | 0.1543 |
| 168 | PW.Jarvis | England | -0.0559 | -0.2836 | 0.1684 |
| 169 | BP.Julian | Australia | -0.0585 | -0.2978 | 0.1769 |
| 170 | RMH.Binny | India | -0.0587 | -0.2622 | 0.1415 |
| 171 | Tapash.Baisya | Bangladesh | -0.0606 | -0.2854 | 0.1566 |
| 172 | ML.Su'a | New Zealand | -0.0616 | -0.2773 | 0.1452 |
| 173 | KSC.de.Silva | SriLanka | -0.0630 | -0.3200 | 0.1753 |
| 174 | MS.Kasprowicz | Australia | -0.0631 | -0.2313 | 0.1022 |
| 175 | PJW.Allott | England | -0.0646 | -0.2866 | 0.1502 |
| 176 | MM.Patel | India | -0.0646 | -0.2813 | 0.1365 |
| 177 | Tahir.Naqqash | Pakistan | -0.0650 | -0.2855 | 0.1465 |
| 178 | BS.Sandhu | India | -0.0661 | -0.3180 | 0.1711 |
| 179 | DJ.Bravo | West Indies | -0.0666 | -0.2490 | 0.1067 |
| 180 | M.Mbangwa | Zimbabwe | -0.0673 | -0.2829 | 0.1415 |
| 181 | AJ.Hall | South Africa | -0.0704 | -0.2754 | 0.1271 |

| | | | | | |
|---|---|---|---|---|---|
| 182 | DR.Gilbert | Australia | -0.0705 | -0.3146 | 0.1571 |
| 183 | A.Flintoff | England | -0.0717 | -0.2166 | 0.0655 |
| 184 | Mohammad.Sharif | Bangladesh | -0.0721 | -0.3232 | 0.1659 |
| 185 | CC.Lewis | England | -0.0721 | -0.2457 | 0.0988 |
| 186 | TL.Best | West Indies | -0.0724 | -0.2713 | 0.1185 |
| 187 | NW.Bracken | Australia | -0.0734 | -0.3212 | 0.1658 |
| 188 | EA.Brandes | Zimbabwe | -0.0738 | -0.2986 | 0.1444 |
| 189 | CJ.Anderson | New Zealand | -0.0739 | -0.3132 | 0.1592 |
| 190 | Mushfiqur.Rahman | Bangladesh | -0.0756 | -0.3167 | 0.1553 |
| 191 | NavedNAulNAHasan | Pakistan | -0.0759 | -0.3118 | 0.1531 |
| 192 | KMDN.Kulasekara | SriLanka | -0.0767 | -0.2799 | 0.1167 |
| 193 | AlNAAmin.Hossain | Bangladesh | -0.0774 | -0.3384 | 0.1679 |
| 194 | PJ.Martin | England | -0.0782 | -0.3149 | 0.1532 |
| 195 | MC.Snedden | New Zealand | -0.0801 | -0.2815 | 0.1197 |
| 196 | CR.Matthews | South Africa | -0.0835 | -0.2858 | 0.1196 |
| 197 | DR.Pringle | England | -0.0914 | -0.2803 | 0.0980 |
| 198 | GI.Allott | New Zealand | -0.0921 | -0.3292 | 0.1313 |
| 199 | BJ.Arnel | New Zealand | -0.0925 | -0.3309 | 0.1421 |
| 200 | C.White | England | -0.0925 | -0.2918 | 0.0927 |
| 201 | GP.Wickramasinghe | SriLanka | -0.0950 | -0.2829 | 0.0870 |
| 202 | AD.Mullally | England | -0.0995 | -0.2930 | 0.0855 |
| 203 | Mohammad.Akram | Pakistan | -0.0995 | -0.3381 | 0.1260 |
| 204 | Aaqib.Javed | Pakistan | -0.1003 | -0.3024 | 0.0973 |
| 205 | Manjural.Islam | Bangladesh | -0.1003 | -0.3235 | 0.1144 |
| 206 | E.Chigumbura | Zimbabwe | -0.1009 | -0.3388 | 0.1192 |
| 207 | Khaled.Mahmud | Bangladesh | -0.1023 | -0.3443 | 0.1278 |
| 208 | SP.O'Donnell | Australia | -0.1041 | -0.3602 | 0.1320 |
| 209 | AB.Agarkar | India | -0.1088 | -0.3072 | 0.0825 |
| 210 | DJ.Capel | England | -0.1105 | -0.3441 | 0.1102 |
| 211 | W.Watson | New Zealand | -0.1129 | -0.3275 | 0.0949 |
| 212 | DJG.Sammy | West Indies | -0.1212 | -0.3033 | 0.0536 |
| 213 | C.Pringle | New Zealand | -0.1230 | -0.3569 | 0.0917 |
| 214 | M.Prabhakar | India | -0.1262 | -0.3019 | 0.0467 |
| 215 | Mohammad.Sami | Pakistan | -0.1349 | -0.3105 | 0.0396 |
| 216 | Abdul.Razzaq | Pakistan | -0.1362 | -0.3162 | 0.0432 |
| 217 | ML.Nkala | Zimbabwe | -0.1408 | -0.3879 | 0.0851 |
| 218 | DBL.Powell | West Indies | -0.1421 | -0.3150 | 0.0285 |
| 219 | EAE.Baptiste | West Indies | -0.1462 | -0.3842 | 0.0702 |
| 220 | Azhar.Mahmood | Pakistan | -0.1464 | -0.3584 | 0.0560 |
| 221 | JDP.Oram | New Zealand | -0.1909 | -0.3925 | 0.0021 |
| 222 | BM.McMillan | South Africa | -0.2107 | -0.3946 | -0.0384 |
| 223 | MF.Maharoof | SriLanka | -0.2114 | -0.4241 | -0.0082 |
| 224 | JH.Kallis | South Africa | -0.2213 | -0.3482 | -0.0952 |
| 225 | UC.Hathurusingha | SriLanka | -0.2257 | -0.4702 | -0.0037 |
| 226 | L.Klusener | South Africa | -0.2761 | -0.4594 | -0.1057 |
| 227 | SR.Watson | Australia | -0.3459 | -0.5383 | -0.1706 |

**Table S7: Rankings for the fast bowlers of modern era (post-1973) Test cricket on the basis of their economy rate.** Estimates represent the player-specific intercepts extracted from the multivariate response models that controlled for various confounding effects. Players that were still thought to be playing Test cricket on or around June 2019 are not included.

| Rank | Player Name | Country | Economy | Econ_lower95CI | Econ_upper95CI |
|---|---|---|---|---|---|
| 1 | SM.Pollock | South Africa | -0.6161 | -0.8110 | -0.4098 |
| 2 | CEL.Ambrose | West Indies | -0.5460 | -0.7460 | -0.3444 |
| 3 | JDP.Oram | New Zealand | -0.5439 | -0.8472 | -0.2584 |
| 4 | M.Mbangwa | Zimbabwe | -0.5265 | -0.9210 | -0.1337 |
| 5 | P.Kumar | India | -0.5103 | -0.9855 | -0.0350 |
| 6 | UC.Hathurusingha | Sri Lanka | -0.4762 | -0.8448 | -0.1234 |
| 7 | CD.Collymore | West Indies | -0.4698 | -0.7607 | -0.1764 |
| 8 | Wasim.Akram | Pakistan | -0.4299 | -0.6317 | -0.2291 |
| 9 | W.Watson | New Zealand | -0.4257 | -0.8127 | -0.0508 |
| 10 | JR.Hammond | Australia | -0.4239 | -0.9611 | 0.0787 |
| 11 | GD.McGrath | Australia | -0.4231 | -0.6074 | -0.2313 |
| 12 | HH.Streak | Zimbabwe | -0.4103 | -0.6601 | -0.1611 |
| 13 | WPUJC.Vaas | Sri Lanka | -0.4015 | -0.6138 | -0.2018 |
| 14 | WKM.Benjamin | West Indies | -0.3938 | -0.7319 | -0.0483 |
| 15 | CA.Walsh | West Indies | -0.3885 | -0.5696 | -0.1964 |
| 16 | CR.Matthews | South Africa | -0.3852 | -0.7522 | -0.0243 |
| 17 | BW.Hilfenhaus | Australia | -0.3780 | -0.6924 | -0.0620 |
| 18 | Mohammad.Asif | Pakistan | -0.3693 | -0.7128 | -0.0282 |
| 19 | DJ.Nash | New Zealand | -0.3634 | -0.6648 | -0.0567 |
| 20 | IR.Bishop | West Indies | -0.3613 | -0.6267 | -0.1021 |
| 21 | C.Pringle | New Zealand | -0.3348 | -0.7276 | 0.0792 |
| 22 | PJ.Martin | England | -0.3336 | -0.7990 | 0.1193 |
| 23 | BA.Reid | Australia | -0.3317 | -0.6621 | -0.0105 |
| 24 | J.Garner | West Indies | -0.3248 | -0.5802 | -0.0702 |
| 25 | KJ.Abbott | South Africa | -0.3217 | -0.7461 | 0.1061 |
| 26 | NA.Foster | England | -0.3215 | -0.6450 | -0.0031 |
| 27 | ARC.Fraser | England | -0.3167 | -0.5803 | -0.0598 |
| 28 | T.Panyangara | Zimbabwe | -0.3122 | -0.7532 | 0.1305 |
| 29 | Shabbir.Ahmed | Pakistan | -0.3072 | -0.7212 | 0.0971 |
| 30 | AD.Mullally | England | -0.3029 | -0.6524 | 0.0380 |
| 31 | MA.Ealham | England | -0.3008 | -0.7841 | 0.1571 |
| 32 | SR.Clark | Australia | -0.2991 | -0.6084 | 0.0221 |
| 33 | N.Kapil.Dev | India | -0.2833 | -0.4839 | -0.0854 |
| 34 | Harbhajan.Singh | India | -0.2807 | -0.4782 | -0.0835 |
| 35 | DT.Hondo | Zimbabwe | -0.2799 | -0.7353 | 0.1619 |
| 36 | ST.Clarke | West Indies | -0.2793 | -0.7124 | 0.1289 |
| 37 | AIC.Dodemaide | Australia | -0.2768 | -0.7062 | 0.1468 |
| 38 | Abdul.Razzaq | Pakistan | -0.2756 | -0.5585 | -0.0129 |
| 39 | PAJ.DeFreitas | England | -0.2746 | -0.5327 | -0.0131 |
| 40 | CE.Cuffy | West Indies | -0.2677 | -0.6760 | 0.1000 |
| 41 | A.Flintoff | England | -0.2655 | -0.4908 | -0.0485 |

| | | | | | |
|---|---|---|---|---|---|
| 42 | SP.O'Donnell | Australia | -0.2642 | -0.7406 | 0.2198 |
| 43 | GC.Small | England | -0.2624 | -0.6146 | 0.1008 |
| 44 | JH.Kallis | South Africa | -0.2576 | -0.4382 | -0.0729 |
| 45 | MD.Marshall | West Indies | -0.2550 | -0.4748 | -0.0299 |
| 46 | PR.Reiffel | Australia | -0.2542 | -0.5288 | 0.0323 |
| 47 | EAE.Baptiste | West Indies | -0.2492 | -0.6513 | 0.1574 |
| 48 | JN.Gillespie | Australia | -0.2374 | -0.4601 | -0.0094 |
| 49 | SR.Watson | Australia | -0.2273 | -0.4811 | 0.0298 |
| 50 | RJ.Sidebottom | England | -0.2231 | -0.5763 | 0.1185 |
| 51 | AH.Gray | West Indies | -0.2205 | -0.7070 | 0.2554 |
| 52 | DNT.Zoysa | Sri Lanka | -0.2180 | -0.5292 | 0.0918 |
| 53 | NW.Bracken | Australia | -0.2155 | -0.7025 | 0.2746 |
| 54 | Aaqib.Javed | Pakistan | -0.2143 | -0.5580 | 0.1259 |
| 55 | TT.Bresnan | England | -0.2102 | -0.5580 | 0.1277 |
| 56 | DR.Tuffey | New Zealand | -0.2095 | -0.5322 | 0.1127 |
| 57 | R.Rampaul | West Indies | -0.2087 | -0.5610 | 0.1644 |
| 58 | CJ.Anderson | New Zealand | -0.2007 | -0.6366 | 0.2248 |
| 59 | PJW.Allott | England | -0.1975 | -0.6003 | 0.2089 |
| 60 | CG.Rackemann | Australia | -0.1935 | -0.6049 | 0.2067 |
| 61 | JR.Ratnayeke | Sri Lanka | -0.1927 | -0.5803 | 0.1742 |
| 62 | Manjural.Islam | Bangladesh | -0.1834 | -0.5830 | 0.2117 |
| 63 | CJ.Jordan | England | -0.1807 | -0.6377 | 0.2672 |
| 64 | Azhar.Mahmood | Pakistan | -0.1760 | -0.5124 | 0.1693 |
| 65 | A.Nel | South Africa | -0.1725 | -0.4574 | 0.1097 |
| 66 | SB.O'Connor | New Zealand | -0.1670 | -0.5237 | 0.1830 |
| 67 | PS.de.Villiers | South Africa | -0.1661 | -0.5307 | 0.1940 |
| 68 | CEH.Croft | West Indies | -0.1650 | -0.4741 | 0.1450 |
| 69 | L.Klusener | South Africa | -0.1633 | -0.4195 | 0.1109 |
| 70 | MM.Patel | India | -0.1613 | -0.5412 | 0.2182 |
| 71 | DJG.Sammy | West Indies | -0.1601 | -0.4433 | 0.1285 |
| 72 | KMDN.Kulasekara | Sri Lanka | -0.1567 | -0.4869 | 0.1816 |
| 73 | AtaNAurNARehman | Pakistan | -0.1555 | -0.5631 | 0.2357 |
| 74 | A.Nehra | India | -0.1538 | -0.5199 | 0.2058 |
| 75 | CT.Tremlett | England | -0.1494 | -0.5500 | 0.2544 |
| 76 | GP.Wickramasinghe | Sri Lanka | -0.1488 | -0.4424 | 0.1408 |
| 77 | WW.Daniel | West Indies | -0.1434 | -0.5750 | 0.2748 |
| 78 | BKV.Prasad | India | -0.1325 | -0.4297 | 0.1711 |
| 79 | SB.Doull | New Zealand | -0.1322 | -0.4345 | 0.1646 |
| 80 | MC.Snedden | New Zealand | -0.1310 | -0.4832 | 0.2189 |
| 81 | ML.Nkala | Zimbabwe | -0.1270 | -0.5689 | 0.3242 |
| 82 | DJ.Bravo | West Indies | -0.1242 | -0.4103 | 0.1565 |
| 83 | BN.Schultz | South Africa | -0.1185 | -0.5478 | 0.3316 |
| 84 | CC.Lewis | England | -0.1169 | -0.4105 | 0.1789 |
| 85 | Mashrafe.Mortaza | Bangladesh | -0.1150 | -0.4474 | 0.2075 |
| 86 | A.Kuruvilla | India | -0.1143 | -0.5453 | 0.3102 |
| 87 | VC.Drakes | West Indies | -0.1087 | -0.5113 | 0.2864 |
| 88 | GF.Lawson | Australia | -0.1086 | -0.3785 | 0.1650 |

| | | | | | |
|---|---|---|---|---|---|
| 89 | DW.Fleming | Australia | -0.1064 | -0.4330 | 0.2297 |
| 90 | SJ.Harmison | England | -0.1040 | -0.3337 | 0.1395 |
| 91 | J.Srinath | India | -0.1025 | -0.3310 | 0.1320 |
| 92 | CR.Miller | Australia | -0.1009 | -0.4547 | 0.2458 |
| 93 | RJ.Harris | Australia | -0.0999 | -0.4135 | 0.2042 |
| 94 | PM.Siddle | Australia | -0.0961 | -0.3337 | 0.1467 |
| 95 | Khaled.Mahmud | Bangladesh | -0.0950 | -0.5111 | 0.3175 |
| 96 | M.Morkel | South Africa | -0.0908 | -0.3155 | 0.1491 |
| 97 | GR.Dilley | England | -0.0828 | -0.3773 | 0.2164 |
| 98 | DR.Pringle | England | -0.0757 | -0.3791 | 0.2194 |
| 99 | IK.Pathan | India | -0.0592 | -0.3646 | 0.2448 |
| 100 | RJ.Ratnayake | Sri Lanka | -0.0573 | -0.4168 | 0.2940 |
| 101 | Tahir.Naqqash | Pakistan | -0.0478 | -0.4551 | 0.3478 |
| 102 | M.Dillon | West Indies | -0.0450 | -0.3280 | 0.2401 |
| 103 | RM.Ellison | England | -0.0423 | -0.4759 | 0.3928 |
| 104 | DR.Gilbert | Australia | -0.0385 | -0.5098 | 0.4223 |
| 105 | CJ.McDermott | Australia | -0.0347 | -0.2525 | 0.1953 |
| 106 | DE.Bollinger | Australia | -0.0269 | -0.4365 | 0.3818 |
| 107 | DJ.Terbrugge | South Africa | -0.0246 | -0.4859 | 0.4420 |
| 108 | AJ.Hall | South Africa | -0.0206 | -0.3673 | 0.3062 |
| 109 | BM.McMillan | South Africa | -0.0164 | -0.3194 | 0.2750 |
| 110 | AB.Agarkar | India | -0.0158 | -0.3332 | 0.3008 |
| 111 | TM.Alderman | Australia | -0.0155 | -0.2894 | 0.2581 |
| 112 | BT.Watambwa | Zimbabwe | -0.0138 | -0.4869 | 0.4691 |
| 113 | Saleem.Jaffar | Pakistan | -0.0138 | -0.4148 | 0.3826 |
| 114 | SP.Jones | England | -0.0124 | -0.3599 | 0.3353 |
| 115 | BP.Julian | Australia | -0.0084 | -0.4594 | 0.4468 |
| 116 | ML.Su'a | New Zealand | -0.0045 | -0.4099 | 0.3826 |
| 117 | FA.Rose | West Indies | -0.0037 | -0.3670 | 0.3442 |
| 118 | M.Prabhakar | India | -0.0025 | -0.2793 | 0.2796 |
| 119 | Azeem.Hafeez | Pakistan | 0.0008 | -0.3855 | 0.3989 |
| 120 | MG.Hughes | Australia | 0.0061 | -0.2333 | 0.2550 |
| 121 | WW.Davis | West Indies | 0.0086 | -0.3585 | 0.3672 |
| 122 | GF.Labrooy | Sri Lanka | 0.0110 | -0.4367 | 0.4437 |
| 123 | MF.Maharoof | Sri Lanka | 0.0119 | -0.3392 | 0.3509 |
| 124 | M.Ntini | South Africa | 0.0128 | -0.1868 | 0.2199 |
| 125 | MR.Whitney | Australia | 0.0131 | -0.3885 | 0.3996 |
| 126 | AR.Caddick | England | 0.0192 | -0.2137 | 0.2463 |
| 127 | Waqar.Younis | Pakistan | 0.0195 | -0.1950 | 0.2315 |
| 128 | JJC.Lawson | West Indies | 0.0226 | -0.3462 | 0.3951 |
| 129 | DK.Liyanage | Sri Lanka | 0.0229 | -0.4197 | 0.4578 |
| 130 | NAM.McLean | West Indies | 0.0272 | -0.3094 | 0.3856 |
| 131 | PW.Jarvis | England | 0.0347 | -0.3832 | 0.4621 |
| 132 | LS.Pascoe | Australia | 0.0445 | -0.3411 | 0.4279 |
| 133 | CK.Langeveldt | South Africa | 0.0493 | -0.4413 | 0.5266 |
| 134 | E.Chigumbura | Zimbabwe | 0.0633 | -0.3388 | 0.4696 |
| 135 | KD.Mills | New Zealand | 0.0648 | -0.2987 | 0.4278 |

| | | | | | |
|---|---|---|---|---|---|
| 136 | VRV.Singh | India | 0.0676 | -0.4136 | 0.5489 |
| 137 | DW.Headley | England | 0.0716 | -0.3093 | 0.4663 |
| 138 | BS.Sandhu | India | 0.0749 | -0.4157 | 0.5688 |
| 139 | AlNAAmin.Hossain | Bangladesh | 0.0753 | -0.4326 | 0.5779 |
| 140 | AA.Donald | South Africa | 0.0757 | -0.1606 | 0.3170 |
| 141 | Shahid.Nazir | Pakistan | 0.0784 | -0.2917 | 0.4500 |
| 142 | RD.King | West Indies | 0.0885 | -0.2556 | 0.4312 |
| 143 | BJ.Arnel | New Zealand | 0.0998 | -0.3696 | 0.5829 |
| 144 | EA.Brandes | Zimbabwe | 0.1055 | -0.3173 | 0.5338 |
| 145 | AJ.Tudor | England | 0.1075 | -0.3034 | 0.5241 |
| 146 | Mohammad.Akram | Pakistan | 0.1144 | -0.3270 | 0.5708 |
| 147 | Junaid.Khan | Pakistan | 0.1170 | -0.2907 | 0.4989 |
| 148 | Shoaib.Akhtar | Pakistan | 0.1189 | -0.1470 | 0.3829 |
| 149 | M.Hayward | South Africa | 0.1199 | -0.2360 | 0.4701 |
| 150 | N.Phillip | West Indies | 0.1265 | -0.3147 | 0.5790 |
| 151 | RMS.Eranga | Sri Lanka | 0.1293 | -0.2520 | 0.5058 |
| 152 | IE.O'Brien | New Zealand | 0.1314 | -0.1924 | 0.4827 |
| 153 | JL.Pattinson | Australia | 0.1317 | -0.2370 | 0.5069 |
| 154 | TCB.Fernando | Sri Lanka | 0.1341 | -0.3076 | 0.5816 |
| 155 | CPH.Ramanayake | Sri Lanka | 0.1350 | -0.2265 | 0.5073 |
| 156 | TJ.Friend | Zimbabwe | 0.1365 | -0.2702 | 0.5446 |
| 157 | MJ.Hoggard | England | 0.1374 | -0.0913 | 0.3616 |
| 158 | Aamer.Nazir | Pakistan | 0.1393 | -0.3086 | 0.6104 |
| 159 | S.Sreesanth | India | 0.1414 | -0.1572 | 0.4404 |
| 160 | KSC.de.Silva | Sri Lanka | 0.1418 | -0.3023 | 0.5925 |
| 161 | PT.Collins | West Indies | 0.1456 | -0.1531 | 0.4287 |
| 162 | PDRL.Perera | Sri Lanka | 0.1606 | -0.2910 | 0.6160 |
| 163 | IG.Butler | New Zealand | 0.1699 | -0.2739 | 0.6336 |
| 164 | L.Balaji | India | 0.1718 | -0.2664 | 0.6089 |
| 165 | DBL.Powell | West Indies | 0.1760 | -0.0966 | 0.4639 |
| 166 | KCG.Benjamin | West Indies | 0.1762 | -0.1270 | 0.4846 |
| 167 | Z.Khan | India | 0.1900 | -0.0192 | 0.3974 |
| 168 | JEC.Franklin | New Zealand | 0.1982 | -0.1108 | 0.4976 |
| 169 | MI.Black | West Indies | 0.2030 | -0.2855 | 0.6913 |
| 170 | MB.Owens | New Zealand | 0.2036 | -0.2318 | 0.6600 |
| 171 | ST.Finn | England | 0.2083 | -0.1041 | 0.5242 |
| 172 | Mohammad.Sami | Pakistan | 0.2106 | -0.0840 | 0.4954 |
| 173 | DAJ.Bracewell | New Zealand | 0.2117 | -0.1132 | 0.5442 |
| 174 | SE.Bond | New Zealand | 0.2133 | -0.1461 | 0.5852 |
| 175 | GI.Allott | New Zealand | 0.2178 | -0.2084 | 0.6676 |
| 176 | CS.Martin | New Zealand | 0.2180 | -0.0072 | 0.4475 |
| 177 | D.Gough | England | 0.2190 | -0.0196 | 0.4604 |
| 178 | KR.Pushpakumara | Sri Lanka | 0.2218 | -0.1127 | 0.5581 |
| 179 | MG.Johnson | Australia | 0.2254 | -0.0011 | 0.4476 |
| 180 | RMH.Binny | India | 0.2296 | -0.1207 | 0.5741 |
| 181 | Umar.Gul | Pakistan | 0.2298 | -0.0448 | 0.5050 |
| 182 | C.White | England | 0.2305 | -0.0751 | 0.5410 |

| | | | | | |
|---|---|---|---|---|---|
| 183 | DV.Lawrence | England | 0.2392 | -0.2342 | 0.7399 |
| 184 | Mohsin.Kamal | Pakistan | 0.2480 | -0.2053 | 0.7136 |
| 185 | DJ.Capel | England | 0.2568 | -0.1313 | 0.6361 |
| 186 | RP.Singh | India | 0.2587 | -0.1233 | 0.6417 |
| 187 | T.Thushara | Sri Lanka | 0.2669 | -0.1683 | 0.6974 |
| 188 | DK.Morrison | New Zealand | 0.2706 | 0.0033 | 0.5327 |
| 189 | DG.Cork | England | 0.2729 | -0.0029 | 0.5467 |
| 190 | M.Zondeki | South Africa | 0.2933 | -0.1857 | 0.7785 |
| 191 | CEW.Silverwood | England | 0.2951 | -0.2042 | 0.7822 |
| 192 | Mushfiqur.Rahman | Bangladesh | 0.2972 | -0.1550 | 0.7541 |
| 193 | LE.Plunkett | England | 0.2977 | -0.0700 | 0.6858 |
| 194 | Wahab.Riaz | Pakistan | 0.2999 | -0.0751 | 0.6758 |
| 195 | CL.Cairns | New Zealand | 0.3010 | 0.0719 | 0.5362 |
| 196 | Talha.Jubair | Bangladesh | 0.3074 | -0.1848 | 0.8164 |
| 197 | NG.Cowans | England | 0.3108 | -0.0467 | 0.6835 |
| 198 | G.Onions | England | 0.3138 | -0.1164 | 0.7640 |
| 199 | Robiul.Islam | Bangladesh | 0.3274 | -0.1343 | 0.7835 |
| 200 | UWMBCA.Welegedara | Sri Lanka | 0.3299 | -0.0390 | 0.7026 |
| 201 | MS.Kasprowicz | Australia | 0.3329 | 0.0616 | 0.6082 |
| 202 | Sohail.Khan | Pakistan | 0.3434 | -0.1231 | 0.8230 |
| 203 | C.Sharma | India | 0.3488 | -0.0117 | 0.6891 |
| 204 | JE.Taylor | West Indies | 0.3520 | 0.0786 | 0.6115 |
| 205 | Tapash.Baisya | Bangladesh | 0.3530 | -0.0355 | 0.7341 |
| 206 | AM.Blignaut | Zimbabwe | 0.3570 | -0.0209 | 0.7278 |
| 207 | B.Lee | Australia | 0.3687 | 0.1502 | 0.5852 |
| 208 | Mohammad.Sharif | Bangladesh | 0.3733 | -0.0951 | 0.8564 |
| 209 | ALF.de.Mel | Sri Lanka | 0.3839 | -0.0192 | 0.7704 |
| 210 | A.Sanford | West Indies | 0.4011 | -0.0088 | 0.8158 |
| 211 | DE.Malcolm | England | 0.4063 | 0.1399 | 0.6712 |
| 212 | SI.Mahmood | England | 0.4141 | -0.0308 | 0.8718 |
| 213 | SL.Malinga | Sri Lanka | 0.4200 | 0.1130 | 0.7312 |
| 214 | KTGD.Prasad | Sri Lanka | 0.4401 | 0.0929 | 0.7949 |
| 215 | TL.Best | West Indies | 0.4438 | 0.1159 | 0.7637 |
| 216 | DA.Stirling | New Zealand | 0.4552 | -0.0318 | 0.9728 |
| 217 | NavedNAulNAHasan | Pakistan | 0.4631 | 0.0392 | 0.8964 |
| 218 | BP.Patterson | West Indies | 0.4717 | 0.1786 | 0.7791 |
| 219 | HK.Olonga | Zimbabwe | 0.4816 | 0.1565 | 0.8053 |
| 220 | CRD.Fernando | Sri Lanka | 0.4924 | 0.1949 | 0.7743 |
| 221 | DH.Brain | Zimbabwe | 0.5223 | 0.0992 | 0.9539 |
| 222 | FH.Edwards | West Indies | 0.5381 | 0.3063 | 0.7814 |
| 223 | VR.Aaron | India | 0.5463 | 0.1010 | 1.0131 |
| 224 | AJ.Bichel | Australia | 0.5731 | 0.2214 | 0.9248 |
| 225 | JDS.Neesham | New Zealand | 0.5779 | 0.1260 | 1.0288 |
| 226 | CEL.Stuart | West Indies | 0.5809 | 0.1228 | 1.0495 |
| 227 | Shahadat.Hossain | Bangladesh | 0.7362 | 0.4133 | 1.0519 |

**Table S8: Rankings for the slow bowlers of modern era (post-1973) Test cricket on the basis of their wicket-taking ability.** Estimates represent the player-specific intercepts extracted from the multivariate response models that controlled for various confounding effects, including the number of overs bowled per innings. Players that were still thought to be playing Test cricket on or around June 2019 are not included.

| Rank | PlayerName | Country | Style | Wickets | Wickets_lower95CI | Wickets_Upper95CI |
|---|---|---|---|---|---|---|
| 1 | M.Muralitharan | Sri Lanka | Spin | 0.4779 | 0.3351 | 0.6138 |
| 2 | SK.Warne | Australia | Spin | 0.4390 | 0.2917 | 0.5736 |
| 3 | SCG.MacGill | Australia | Spin | 0.3591 | 0.1698 | 0.5418 |
| 4 | A.Kumble | India | Spin | 0.3464 | 0.2080 | 0.4908 |
| 5 | GP.Swann | England | Spin | 0.3319 | 0.1413 | 0.5265 |
| 6 | Danish.Kaneria | Pakistan | Spin | 0.3004 | 0.1182 | 0.4834 |
| 7 | Saeed.Ajmal | Pakistan | Spin | 0.2911 | 0.0625 | 0.5225 |
| 8 | D.Ramnarine | West Indies | Spin | 0.2566 | -0.0454 | 0.5504 |
| 9 | ND.Hirwani | India | Spin | 0.2552 | -0.0068 | 0.5090 |
| 10 | Saqlain.Mushtaq | Pakistan | Spin | 0.2408 | 0.0530 | 0.4245 |
| 11 | Mushtaq.Ahmed | Pakistan | Spin | 0.2250 | 0.0210 | 0.4198 |
| 12 | PP.Ojha | India | Spin | 0.2247 | -0.0128 | 0.4586 |
| 13 | Abdur.Rehman | Pakistan | Spin | 0.2178 | -0.0551 | 0.4857 |
| 14 | PR.Adams | South Africa | Spin | 0.2072 | 0.0020 | 0.4137 |
| 15 | S.Shillingford | West Indies | Spin | 0.2061 | -0.0572 | 0.4734 |
| 16 | NM.Hauritz | Australia | Spin | 0.2052 | -0.0508 | 0.4690 |
| 17 | A.Mishra | India | Spin | 0.1956 | -0.0493 | 0.4467 |
| 18 | NGB.Cook | England | Spin | 0.1797 | -0.1059 | 0.4777 |
| 19 | DL.Piedt | South Africa | Spin | 0.1739 | -0.1682 | 0.5289 |
| 20 | KPJ.Warnaweera | Sri Lanka | Spin | 0.1738 | -0.1501 | 0.4903 |
| 21 | DS.de.Silva | Sri Lanka | Spin | 0.1724 | -0.1393 | 0.4829 |
| 22 | KJ.Silva | Sri Lanka | Spin | 0.1642 | -0.1803 | 0.5116 |
| 23 | AJ.Traicos | South Africa | Spin | 0.1582 | -0.1684 | 0.4801 |
| 24 | JG.Bracewell | New Zealand | Spin | 0.1394 | -0.0875 | 0.3720 |
| 25 | MS.Panesar | England | Spin | 0.1369 | -0.0591 | 0.3389 |
| 26 | Mohammad.Rafique | Bangladesh | Spin | 0.1331 | -0.0972 | 0.3740 |
| 27 | BAW.Mendis | Sri Lanka | Spin | 0.1304 | -0.1245 | 0.3809 |
| 28 | AG.Huckle | Zimbabwe | Spin | 0.1273 | -0.2084 | 0.4616 |
| 29 | SJ.Benn | West Indies | Spin | 0.1251 | -0.1043 | 0.3593 |
| 30 | Jubair.Hossain | Bangladesh | Spin | 0.1244 | -0.2625 | 0.5285 |
| 31 | DR.Doshi | India | Spin | 0.1234 | -0.1025 | 0.3403 |
| 32 | S.Randiv | Sri Lanka | Spin | 0.1164 | -0.1936 | 0.4158 |
| 33 | SLV.Raju | India | Spin | 0.1102 | -0.1186 | 0.3346 |
| 34 | GR.Larsen | New Zealand | SlowMedium | 0.1094 | -0.2259 | 0.4318 |
| 35 | Sohag.Gazi | Bangladesh | Spin | 0.1044 | -0.2107 | 0.4321 |
| 36 | RG.Holland | Australia | Spin | 0.1008 | -0.2150 | 0.4138 |
| 37 | MD.Craig | New Zealand | Spin | 0.0998 | -0.2129 | 0.4067 |
| 38 | PHT.Kaushal | Sri Lanka | Spin | 0.0997 | -0.2345 | 0.4365 |
| 39 | TBA.May | Australia | Spin | 0.0963 | -0.1364 | 0.3290 |
| 40 | Arshad.Ayub | India | Spin | 0.0949 | -0.2028 | 0.3792 |

| | | | | | | |
|---|---|---|---|---|---|---|
| 41 | PM.Such | England | Spin | 0.0936 | -0.2179 | 0.3965 |
| 42 | M.Kartik | India | Spin | 0.0867 | -0.2717 | 0.4210 |
| 43 | RA.Harper | West Indies | Spin | 0.0814 | -0.2121 | 0.3628 |
| 44 | TV.Hohns | Australia | Spin | 0.0800 | -0.2905 | 0.4256 |
| 45 | TAM.Siriwardana | Sri Lanka | Spin | 0.0790 | -0.3203 | 0.4678 |
| 46 | Imran.Tahir | South Africa | Spin | 0.0718 | -0.2098 | 0.3563 |
| 47 | Enamul.Haque.jnr | Bangladesh | Spin | 0.0699 | -0.2097 | 0.3635 |
| 48 | NS.Yadav | India | Spin | 0.0609 | -0.1600 | 0.2825 |
| 49 | MRCN.Bandaratilleke | Sri Lanka | Spin | 0.0585 | -0.2765 | 0.3981 |
| 50 | RJ.Peterson | South Africa | Spin | 0.0534 | -0.2419 | 0.3346 |
| 51 | D.Mohammed | West Indies | Spin | 0.0491 | -0.3376 | 0.4299 |
| 52 | TG.Hogan | Australia | Spin | 0.0470 | -0.3112 | 0.4017 |
| 53 | EE.Hemmings | England | Spin | 0.0438 | -0.2357 | 0.3239 |
| 54 | DL.Vettori | New Zealand | Spin | 0.0420 | -0.1100 | 0.1909 |
| 55 | DR.Parry | West Indies | Spin | 0.0407 | -0.3119 | 0.3814 |
| 56 | Shahid.Afridi | Pakistan | Spin | 0.0372 | -0.2409 | 0.3105 |
| 57 | RK.Illingworth | England | Spin | 0.0343 | -0.3180 | 0.3744 |
| 58 | PL.Harris | South Africa | Spin | 0.0319 | -0.1964 | 0.2540 |
| 59 | RW.Price | Zimbabwe | Spin | 0.0314 | -0.2180 | 0.2758 |
| 60 | Arshad.Khan | Pakistan | Spin | 0.0309 | -0.2987 | 0.3555 |
| 61 | PA.Strang | Zimbabwe | Spin | 0.0309 | -0.2259 | 0.2744 |
| 62 | Mohammad.Nazir | Pakistan | Spin | 0.0283 | -0.3031 | 0.3604 |
| 63 | V.Permaul | West Indies | Spin | 0.0254 | -0.3409 | 0.3872 |
| 64 | Maninder.Singh | India | Spin | 0.0242 | -0.2213 | 0.2573 |
| 65 | SB.Joshi | India | Spin | 0.0215 | -0.2603 | 0.2944 |
| 66 | RR.Jumadeen | West Indies | Spin | 0.0171 | -0.3169 | 0.3392 |
| 67 | AF.Giles | England | Spin | 0.0030 | -0.2015 | 0.2074 |
| 68 | Tauseef.Ahmed | Pakistan | Spin | 0.0017 | -0.2415 | 0.2304 |
| 69 | VJ.Marks | England | Spin | -0.0054 | -0.3991 | 0.3861 |
| 70 | PCR.Tufnell | England | Spin | -0.0067 | -0.2245 | 0.2080 |
| 71 | PI.Pocock | England | Spin | -0.0104 | -0.3073 | 0.2790 |
| 72 | SD.Anurasiri | Sri Lanka | Spin | -0.0131 | -0.3056 | 0.2712 |
| 73 | SP.Narine | West Indies | Spin | -0.0135 | -0.3612 | 0.3335 |
| 74 | L.Sivaramakrishnan | India | Spin | -0.0192 | -0.3540 | 0.3178 |
| 75 | OAC.Banks | West Indies | Spin | -0.0196 | -0.3392 | 0.3032 |
| 76 | JS.Patel | New Zealand | Spin | -0.0212 | -0.2759 | 0.2239 |
| 77 | RDB.Croft | England | Spin | -0.0222 | -0.2907 | 0.2290 |
| 78 | PL.Taylor | Australia | Spin | -0.0231 | -0.3395 | 0.2818 |
| 79 | MN.Hart | New Zealand | Spin | -0.0308 | -0.3568 | 0.2807 |
| 80 | N.Boje | South Africa | Spin | -0.0315 | -0.2568 | 0.1972 |
| 81 | DN.Patel | New Zealand | Spin | -0.0337 | -0.2822 | 0.2026 |
| 82 | HMCM.Bandara | Sri Lanka | Spin | -0.0361 | -0.4082 | 0.3092 |
| 83 | CW.Henderson | South Africa | Spin | -0.0408 | -0.3877 | 0.2815 |
| 84 | RKJ.Dawson | England | Spin | -0.0426 | -0.4249 | 0.3349 |
| 85 | PL.Symcox | South Africa | Spin | -0.0503 | -0.3536 | 0.2366 |
| 86 | CG.Butts | West Indies | Spin | -0.0517 | -0.4382 | 0.3214 |
| 87 | GB.Hogg | Australia | Spin | -0.0559 | -0.4220 | 0.3005 |

| | | | | | | |
|---|---|---|---|---|---|---|
| 88 | RK.Chauhan | India | Spin | -0.0594 | -0.3335 | 0.2060 |
| 89 | ET.Willett | West Indies | Spin | -0.0603 | -0.4562 | 0.2962 |
| 90 | HDPK.Dharmasena | Sri Lanka | Spin | -0.0646 | -0.2986 | 0.1696 |
| 91 | UDU.Chandana | Sri Lanka | Spin | -0.0687 | -0.3701 | 0.2205 |
| 92 | NB.Mahwire | Zimbabwe | SlowMedium | -0.0760 | -0.4525 | 0.2815 |
| 93 | RJ.Bright | Australia | Spin | -0.0789 | -0.3647 | 0.1947 |
| 94 | N.Deonarine | West Indies | Spin | -0.0801 | -0.4078 | 0.2403 |
| 95 | Enamul.Haque | Bangladesh | Spin | -0.0868 | -0.4346 | 0.2637 |
| 96 | EJ.Gray | New Zealand | Spin | -0.0896 | -0.4398 | 0.2608 |
| 97 | BC.Strang | Zimbabwe | SlowMedium | -0.0968 | -0.3559 | 0.1636 |
| 98 | PJ.Wiseman | New Zealand | Spin | -0.1176 | -0.3642 | 0.1260 |
| 99 | SR.Patel | England | Spin | -0.1205 | -0.5329 | 0.2752 |
| 100 | Akram.Raza | Pakistan | Spin | -0.1232 | -0.4771 | 0.2286 |
| 101 | BA.Murphy | Zimbabwe | Spin | -0.1345 | -0.4798 | 0.1839 |
| 102 | RJ.Shastri | India | Spin | -0.1575 | -0.3515 | 0.0390 |
| 103 | RS.Kalpage | Sri Lanka | Spin | -0.1650 | -0.5333 | 0.1835 |
| 104 | GJ.Batty | England | Spin | -0.1671 | -0.5298 | 0.1846 |
| 105 | Naimur.Rahman | Bangladesh | Spin | -0.1674 | -0.5614 | 0.2102 |
| 106 | SA.Thomson | New Zealand | Spin | -0.1792 | -0.5203 | 0.1458 |
| 107 | MM.Patel | England | Spin | -0.1897 | -0.6536 | 0.2602 |
| 108 | GRJ.Matthews | Australia | Spin | -0.1949 | -0.4514 | 0.0550 |
| 109 | Mohammad.Hafeez | Pakistan | Spin | -0.2077 | -0.4904 | 0.0675 |
| 110 | S.Shivnarine | West Indies | Spin | -0.2164 | -0.6484 | 0.1937 |
| 111 | Mehrab.Hossain.jnr | Bangladesh | Spin | -0.2192 | -0.6603 | 0.1960 |
| 112 | RO.Hinds | West Indies | Spin | -0.2262 | -0.5959 | 0.1215 |
| 113 | MLC.Foster | West Indies | Spin | -0.2298 | -0.6292 | 0.1301 |
| 114 | Shuvagata.Hom | Bangladesh | Spin | -0.2308 | -0.6269 | 0.1465 |
| 115 | CS.Cowdrey | England | SlowMedium | -0.2309 | -0.6827 | 0.1986 |
| 116 | A.Symonds | Australia | SlowMedium | -0.2456 | -0.5616 | 0.0508 |
| 117 | ED.Solkar | India | Spin | -0.2509 | -0.6966 | 0.1373 |
| 118 | IDK.Salisbury | England | Spin | -0.2641 | -0.6123 | 0.0611 |
| 119 | CE.Eksteen | South Africa | Spin | -0.2819 | -0.6762 | 0.0742 |
| 120 | KBJ.Azad | India | Spin | -0.2853 | -0.6962 | 0.1090 |
| 121 | CL.Hooper | West Indies | Spin | -0.3145 | -0.5328 | -0.1076 |
| 122 | AR.Whittall | Zimbabwe | Spin | -0.3213 | -0.7178 | 0.0376 |
| 123 | EAR.de.Silva | Sri Lanka | Spin | -0.3379 | -0.7304 | 0.0196 |
| 124 | ST.Jayasuriya | Sri Lanka | Spin | -0.3578 | -0.5769 | -0.1569 |
| 125 | Alok.Kapali | Bangladesh | Spin | -0.3669 | -0.7812 | -0.0031 |
| 126 | CZ.Harris | New Zealand | SlowMedium | -0.3789 | -0.7283 | -0.0627 |
| 127 | Naeem.Islam | Bangladesh | Spin | -0.3887 | -0.8161 | -0.0057 |
| 128 | Mohammad.Ashraful | Bangladesh | Spin | -0.6483 | -1.0037 | -0.3214 |

**Table S9: Rankings for the slow bowlers of modern era (post-1973) Test cricket on the basis of their economy rate.** Estimates represent the player-specific intercepts extracted from the multivariate response models that controlled for various confounding effects. Players that were still thought to be playing Test cricket on or around June 2019 are not included.

| Rank | PlayerName | Country | Style | Economy | Econ_lower95CI | Econ_upper95CI |
|---|---|---|---|---|---|---|
| 1 | GR.Larsen | New Zealand | SlowMedium | -0.5364 | -1.0993 | -0.0163 |
| 2 | RA.Harper | West Indies | Spin | -0.5280 | -0.9756 | -0.1099 |
| 3 | RJ.Bright | Australia | Spin | -0.4722 | -0.8989 | -0.0622 |
| 4 | DR.Doshi | India | Spin | -0.4202 | -0.7842 | -0.0500 |
| 5 | CG.Butts | West Indies | Spin | -0.4191 | -1.0114 | 0.1447 |
| 6 | Mohammad.Rafique | Bangladesh | Spin | -0.4153 | -0.8475 | 0.0054 |
| 7 | A.Symonds | Australia | SlowMedium | -0.4126 | -0.8074 | -0.0320 |
| 8 | PL.Harris | South Africa | Spin | -0.3988 | -0.7602 | -0.0532 |
| 9 | PCR.Tufnell | England | Spin | -0.3905 | -0.7499 | -0.0588 |
| 10 | Arshad.Ayub | India | Spin | -0.3832 | -0.8701 | 0.0899 |
| 11 | M.Kartik | India | Spin | -0.3817 | -0.9042 | 0.1332 |
| 12 | PP.Ojha | India | Spin | -0.3782 | -0.7880 | 0.0365 |
| 13 | Abdur.Rehman | Pakistan | Spin | -0.3760 | -0.8299 | 0.0918 |
| 14 | Arshad.Khan | Pakistan | Spin | -0.3607 | -0.8905 | 0.1468 |
| 15 | Mohammad.Nazir | Pakistan | Spin | -0.3571 | -0.9143 | 0.1647 |
| 16 | Maninder.Singh | India | Spin | -0.3563 | -0.7408 | 0.0322 |
| 17 | M.Muralitharan | Sri Lanka | Spin | -0.3562 | -0.5990 | -0.1008 |
| 18 | Saeed.Ajmal | Pakistan | Spin | -0.3515 | -0.7557 | 0.0390 |
| 19 | SD.Anurasiri | Sri Lanka | Spin | -0.3465 | -0.7848 | 0.0777 |
| 20 | SLV.Raju | India | Spin | -0.3364 | -0.7074 | 0.0310 |
| 21 | D.Ramnarine | West Indies | Spin | -0.3344 | -0.8204 | 0.1414 |
| 22 | TV.Hohns | Australia | Spin | -0.3261 | -0.8883 | 0.2112 |
| 23 | NGB.Cook | England | Spin | -0.3138 | -0.7857 | 0.1422 |
| 24 | PM.Such | England | Spin | -0.3134 | -0.8165 | 0.1621 |
| 25 | EE.Hemmings | England | Spin | -0.2954 | -0.7256 | 0.1418 |
| 26 | Enamul.Haque | Bangladesh | Spin | -0.2928 | -0.8452 | 0.2676 |
| 27 | RK.Chauhan | India | Spin | -0.2585 | -0.6791 | 0.1518 |
| 28 | CL.Hooper | West Indies | Spin | -0.2514 | -0.5408 | 0.0200 |
| 29 | MS.Panesar | England | Spin | -0.2313 | -0.5417 | 0.1004 |
| 30 | KJ.Silva | Sri Lanka | Spin | -0.2283 | -0.8023 | 0.3098 |
| 31 | PI.Pocock | England | Spin | -0.2229 | -0.6922 | 0.2338 |
| 32 | DL.Vettori | New Zealand | Spin | -0.2197 | -0.4649 | 0.0362 |
| 33 | RJ.Shastri | India | Spin | -0.2139 | -0.4970 | 0.0877 |
| 34 | Tauseef.Ahmed | Pakistan | Spin | -0.2095 | -0.5798 | 0.1884 |
| 35 | SJ.Benn | West Indies | Spin | -0.2048 | -0.5877 | 0.1968 |
| 36 | Saqlain.Mushtaq | Pakistan | Spin | -0.1988 | -0.5142 | 0.1221 |
| 37 | CE.Eksteen | South Africa | Spin | -0.1884 | -0.7200 | 0.3390 |
| 38 | HDPK.Dharmasena | Sri Lanka | Spin | -0.1873 | -0.5392 | 0.1537 |
| 39 | TBA.May | Australia | Spin | -0.1806 | -0.5508 | 0.2033 |
| 40 | SK.Warne | Australia | Spin | -0.1649 | -0.4059 | 0.0800 |
| 41 | ET.Willett | West Indies | Spin | -0.1647 | -0.7496 | 0.4151 |

| | | | | | | |
|---|---|---|---|---|---|---|
| 42 | MLC.Foster | West Indies | Spin | -0.1458 | -0.7052 | 0.4002 |
| 43 | RR.Jumadeen | West Indies | Spin | -0.1387 | -0.6524 | 0.3658 |
| 44 | A.Kumble | India | Spin | -0.1329 | -0.3822 | 0.1340 |
| 45 | Enamul.Haque.jnr | Bangladesh | Spin | -0.1304 | -0.6116 | 0.3579 |
| 46 | SB.Joshi | India | Spin | -0.1233 | -0.5746 | 0.3306 |
| 47 | PL.Symcox | South Africa | Spin | -0.1192 | -0.5448 | 0.3217 |
| 48 | BAW.Mendis | Sri Lanka | Spin | -0.1186 | -0.5509 | 0.2893 |
| 49 | KPJ.Warnaweera | Sri Lanka | Spin | -0.1184 | -0.6343 | 0.4128 |
| 50 | EAR.de.Silva | Sri Lanka | Spin | -0.1142 | -0.6413 | 0.4121 |
| 51 | Mohammad.Hafeez | Pakistan | Spin | -0.1130 | -0.4917 | 0.2598 |
| 52 | MRCN.Bandaratilleke | Sri Lanka | Spin | -0.1125 | -0.6488 | 0.3915 |
| 53 | BA.Murphy | Zimbabwe | Spin | -0.1086 | -0.6106 | 0.3903 |
| 54 | JG.Bracewell | New Zealand | Spin | -0.1018 | -0.4777 | 0.2644 |
| 55 | S.Shivnarine | West Indies | Spin | -0.1016 | -0.6950 | 0.5113 |
| 56 | DS.de.Silva | Sri Lanka | Spin | -0.1009 | -0.5881 | 0.4206 |
| 57 | ST.Jayasuriya | Sri Lanka | Spin | -0.0644 | -0.3312 | 0.2053 |
| 58 | TG.Hogan | Australia | Spin | -0.0515 | -0.6494 | 0.5373 |
| 59 | RDB.Croft | England | Spin | -0.0480 | -0.4586 | 0.3597 |
| 60 | SR.Patel | England | Spin | -0.0436 | -0.6504 | 0.5772 |
| 61 | A.Mishra | India | Spin | -0.0427 | -0.4503 | 0.3720 |
| 62 | CW.Henderson | South Africa | Spin | -0.0409 | -0.6038 | 0.5220 |
| 63 | N.Boje | South Africa | Spin | -0.0347 | -0.3776 | 0.3011 |
| 64 | GP.Swann | England | Spin | -0.0219 | -0.3385 | 0.2975 |
| 65 | Akram.Raza | Pakistan | Spin | -0.0214 | -0.5385 | 0.4979 |
| 66 | PA.Strang | Zimbabwe | Spin | -0.0146 | -0.4117 | 0.3876 |
| 67 | RK.Illingworth | England | Spin | -0.0117 | -0.5700 | 0.5477 |
| 68 | AF.Giles | England | Spin | -0.0088 | -0.3493 | 0.3115 |
| 69 | RW.Price | Zimbabwe | Spin | -0.0059 | -0.4259 | 0.4051 |
| 70 | BC.Strang | Zimbabwe | SlowMedium | -0.0001 | -0.3904 | 0.3804 |
| 71 | RS.Kalpage | Sri Lanka | Spin | 0.0008 | -0.5080 | 0.4978 |
| 72 | S.Shillingford | West Indies | Spin | 0.0040 | -0.4377 | 0.4653 |
| 73 | SA.Thomson | New Zealand | Spin | 0.0060 | -0.4351 | 0.4569 |
| 74 | NS.Yadav | India | Spin | 0.0109 | -0.3541 | 0.3761 |
| 75 | GRJ.Matthews | Australia | Spin | 0.0242 | -0.3397 | 0.3938 |
| 76 | DL.Piedt | South Africa | Spin | 0.0299 | -0.5251 | 0.5995 |
| 77 | VJ.Marks | England | Spin | 0.0551 | -0.5233 | 0.6500 |
| 78 | S.Randiv | Sri Lanka | Spin | 0.0600 | -0.4233 | 0.5766 |
| 79 | AJ.Traicos | South Africa | Spin | 0.0621 | -0.4822 | 0.6030 |
| 80 | EJ.Gray | New Zealand | Spin | 0.0634 | -0.4484 | 0.5896 |
| 81 | OAC.Banks | West Indies | Spin | 0.0685 | -0.3994 | 0.5607 |
| 82 | MN.Hart | New Zealand | Spin | 0.0866 | -0.3821 | 0.5475 |
| 83 | PL.Taylor | Australia | Spin | 0.0937 | -0.3651 | 0.5715 |
| 84 | NM.Hauritz | Australia | Spin | 0.0950 | -0.3136 | 0.5016 |
| 85 | RG.Holland | Australia | Spin | 0.1048 | -0.3754 | 0.6024 |
| 86 | Shahid.Afridi | Pakistan | Spin | 0.1126 | -0.2721 | 0.4996 |
| 87 | GJ.Batty | England | Spin | 0.1129 | -0.4113 | 0.6579 |
| 88 | Naeem.Islam | Bangladesh | Spin | 0.1222 | -0.4337 | 0.6517 |

| | | | | | | |
|---|---|---|---|---|---|---|
| 89 | Sohag.Gazi | Bangladesh | Spin | 0.1278 | -0.3689 | 0.6233 |
| 90 | Danish.Kaneria | Pakistan | Spin | 0.1312 | -0.1717 | 0.4567 |
| 91 | ND.Hirwani | India | Spin | 0.1436 | -0.2720 | 0.5674 |
| 92 | V.Permaul | West Indies | Spin | 0.1441 | -0.4117 | 0.6912 |
| 93 | SP.Narine | West Indies | Spin | 0.1561 | -0.4187 | 0.7279 |
| 94 | DR.Parry | West Indies | Spin | 0.1598 | -0.3513 | 0.6826 |
| 95 | Mushtaq.Ahmed | Pakistan | Spin | 0.1853 | -0.1293 | 0.4939 |
| 96 | AR.Whittall | Zimbabwe | Spin | 0.1913 | -0.3657 | 0.7346 |
| 97 | PR.Adams | South Africa | Spin | 0.1967 | -0.1374 | 0.5202 |
| 98 | DN.Patel | New Zealand | Spin | 0.1983 | -0.1685 | 0.5468 |
| 99 | RKJ.Dawson | England | Spin | 0.2090 | -0.3994 | 0.8269 |
| 100 | CZ.Harris | New Zealand | SlowMedium | 0.2123 | -0.2096 | 0.6455 |
| 101 | PHT.Kaushal | Sri Lanka | Spin | 0.2246 | -0.3150 | 0.7901 |
| 102 | ED.Solkar | India | Spin | 0.2261 | -0.3433 | 0.8098 |
| 103 | TAM.Siriwardana | Sri Lanka | Spin | 0.2265 | -0.3826 | 0.8386 |
| 104 | PJ.Wiseman | New Zealand | Spin | 0.2279 | -0.1497 | 0.5841 |
| 105 | SCG.MacGill | Australia | Spin | 0.2378 | -0.0828 | 0.5480 |
| 106 | HMCM.Bandara | Sri Lanka | Spin | 0.2394 | -0.2939 | 0.7840 |
| 107 | RJ.Peterson | South Africa | Spin | 0.2427 | -0.2046 | 0.7012 |
| 108 | JS.Patel | New Zealand | Spin | 0.2449 | -0.1259 | 0.6426 |
| 109 | D.Mohammed | West Indies | Spin | 0.2536 | -0.3324 | 0.8689 |
| 110 | Imran.Tahir | South Africa | Spin | 0.2908 | -0.1647 | 0.7370 |
| 111 | KBJ.Azad | India | Spin | 0.3077 | -0.2539 | 0.8782 |
| 112 | AG.Huckle | Zimbabwe | Spin | 0.3083 | -0.2322 | 0.8529 |
| 113 | Shuvagata.Hom | Bangladesh | Spin | 0.3182 | -0.2540 | 0.9041 |
| 114 | L.Sivaramakrishnan | India | Spin | 0.3306 | -0.1643 | 0.8448 |
| 115 | MM.Patel | England | Spin | 0.3585 | -0.2976 | 1.0663 |
| 116 | UDU.Chandana | Sri Lanka | Spin | 0.3694 | -0.0701 | 0.8001 |
| 117 | RO.Hinds | West Indies | Spin | 0.3696 | -0.1084 | 0.8564 |
| 118 | GB.Hogg | Australia | Spin | 0.3742 | -0.1514 | 0.9243 |
| 119 | N.Deonarine | West Indies | Spin | 0.3889 | -0.0556 | 0.8395 |
| 120 | Alok.Kapali | Bangladesh | Spin | 0.4130 | -0.1024 | 0.9212 |
| 121 | MD.Craig | New Zealand | Spin | 0.4183 | -0.0569 | 0.9072 |
| 122 | IDK.Salisbury | England | Spin | 0.4379 | -0.0089 | 0.9166 |
| 123 | Jubair.Hossain | Bangladesh | Spin | 0.4465 | -0.1488 | 1.0940 |
| 124 | Mehrab.Hossain.jnr | Bangladesh | Spin | 0.5464 | -0.0260 | 1.1628 |
| 125 | Naimur.Rahman | Bangladesh | Spin | 0.7970 | 0.1873 | 1.4100 |
| 126 | CS.Cowdrey | England | SlowMedium | 0.8885 | 0.2716 | 1.5432 |
| 127 | NB.Mahwire | Zimbabwe | SlowMedium | 1.0255 | 0.4719 | 1.6234 |
| 128 | Mohammad.Ashraful | Bangladesh | Spin | 1.7366 | 1.3063 | 2.1417 |
